# Spatial synaptic regularization stabilizes learning across biological and artificial neural networks

**DOI:** 10.64898/2026.06.29.735142

**Authors:** Haoxing Zhu, Yinda Chen, Peng Zhao, Zhiwei Xiong, Hanchuan Peng, Feng Wu, Ruobing Zhang

## Abstract

How the spatial organization of synapses contributes to stable learning remains a fundamental question in neuroscience. Using the H01 electron microscopy connectome of human temporal cortex, we found that dendritic spines clustered morphologically, whereas synaptic weights followed a center-elevated, surround-suppressed arrangement along dendrites. A regularized Hebbian model formalized this spatial signature, showing that strong synapses lower the probability that neighboring synapses reach high-weight states. Translating this principle into Spatial Synaptic Regularization (SSR) reduced forgetting and stabilized learning across diverse artificial networks and tasks, including continual visual learning, large language-model knowledge editing, and parameter-efficient adaptation of vision-language models, by preserving high-rank, low-overlap representations. These findings identify spatial synaptic organization as an unrecognized dimension for stabilizing learning and show that structural connectomics can yield actionable AI methods.

---

Learning requires two seemingly contradictory capacities: the ability to acquire new information and the ability to retain what has already been learned. Synaptic plasticity provides one central substrate for this balance. Classical theories of synaptic plasticity, from Hebbian learning to subsequent extensions such as Oja’s rule and Bienenstock–Cooper–Munro theory, have provided a rich account of how individual synapses strengthen, weaken, or normalize in response to neural activity. These theories are primarily temporal: they specify when synapses change, how strongly they grow, and how weight changes are stabilized over time (*1–9*). Yet synapses are not isolated learning variables. Most of them are physically embedded along dendritic branches, where neighboring synapses share biochemical signals, electrical compartments, and structural resources (*10–16*). Whether spatial relationships among synapses constitute an additional organizing dimension of learning, shaping not only the dynamics of individual synapses but also the rules by which neighboring synapses interact with one another, is unresolved.

Although neighboring synapses are known to interact through local dendritic and heterosynaptic mechanisms, whether these interactions impose selective spatial constraints on the local co-occupation of synaptic weight states remains unknown. This has been difficult to test because such an analysis requires conditions rarely available together: synapse-scale resolution, precise spine positions, and sufficient statistical scale. The H01 human temporal cortex and MICrONS mouse V1 reconstructions now provide this opportunity at the scale of millions of synapses (*17–21*).

To test whether synaptic weight states follow local spatial organization rules, we analyzed synaptic weight organization in the human temporal cortex and the mouse V1 cortex. Neighboring dendritic spines were more morphologically similar than expected by chance, indicating local morphological clustering. Synaptic weights, however, followed a different rule. Large synapses were embedded in a center–surround weight field: immediate local elevation was followed by an intermediate zone of suppression, with recovery at larger distances. This Mexican-hat profile was consistent across cortical layers and cell classes in human temporal cortex, but was absent in mouse V1, revealing a dissociation between spine morphology and synaptic weight organization, which suggests that synaptic weight-state organization is not simply inherited from local spine morphology, but reflects an additional spatial principle.

To formalize this topology, we extended classical Hebbian learning with a spatial interaction term. The model predicts a selective consequence: a strong synapse reduces the probability that a nearby synapse enters a high-weight state, while leaving weak and medium-weight neighbors weakly affected. Permutation analysis confirmed this prediction: high-weight neighbors were selectively suppressed in the suppression zone, whereas weak and medium-weight neighbors remained near their baseline expectation.

Translating the center–surround topology into Spatial Synaptic Regularization (SSR), a regularizer over learned directions in artificial networks, we asked whether this principle retains computational force beyond dendrites. Across matched continual image-classification protocols, SSR reduced forgetting and stabilized plasticity by preserving high-rank, low-overlap representations. Because SSR acts on the geometry of learned directions rather than on task-specific parameters, the same prior can be tested beyond visual classification in language-model knowledge-editing and low-rank adapter probes.

Together, these findings reframe learning stability as a problem of spatial weight-state organization. Classical accounts have focused on the timing, magnitude, and normalization of synaptic weight change. The center-elevation/surround-inhibition topology identified here adds a complementary spatial dimension: stable learning can arise not by freezing synapses or globally suppressing plasticity, but by imposing spatial rules on how synaptic weight states are locally organized and co-occupied. In this view, dendritic geometry functions as a local memory-allocation system — permitting plasticity while preventing high-weight states from saturating the same neighborhood. More broadly, these results suggest that the geometry of synaptic weight states — not only their magnitude or temporal dynamics — is a fundamental organizing principle of learning. By translating this topology into artificial networks, we further show that large-scale connectomics can serve as a source of actionable computational principles, linking nanoscale structural constraints in human cortical synapses to regularization strategies that mitigate catastrophic forgetting in artificial networks.

## Dendritic spine morphology shows local spatial clustering

We first analyzed dendritic spine morphology in the H01 human left temporal-cortex reconstruction and the MICrONS mouse primary visual-cortex reconstruction (*19–21*). After quality filtering and branch assignment, the analysis included 4.34 million H01 spines from 5,340 reconstructed neurons and 1.89 million MICrONS spine-bearing synapses from 1,333 neurons (Fig. 1a,c). Each spine was represented by a z-scored six-dimensional morphology vector describing length, volume, head diameter, neck diameter, head-to-neck diameter ratio, and surface area (Fig. 1b). Spine morphology was locally organized along dendrites. For each spatial window size, we computed the average pairwise Euclidean distance among all spines within the window and compared it with two spine-count-matched controls: global-random samples drawn from the whole neuron and branch-random samples drawn from the same dendritic branch. In H01 pyramidal cells, local spatial-window groups had lower morphological distance than both controls across the tested window range (Fig. 1d). The separation was strongest at small windows and weakened as the window expanded, indicating that local morphological coherence is progressively diluted when more distant spines are included. Branch-random samples were also more compact than global-random samples, indicating broader branch-level structure. Thus, dendritic spine morphology showed a nested spatial organization: local neighborhoods were more coherent than branch-matched random groups, which were in turn more coherent than global random groups. The same ordering was observed across H01 pyramidal neurons, interneurons, spiny stellate cells, and excitatory spiny neurons (Fig. 1e), and was also recovered in MICrONS pyramidal neurons and interneurons (Fig. 1f). These results show that spatially neighboring spines share local morphological microenvironments beyond branch identity.

**Figure 1:**
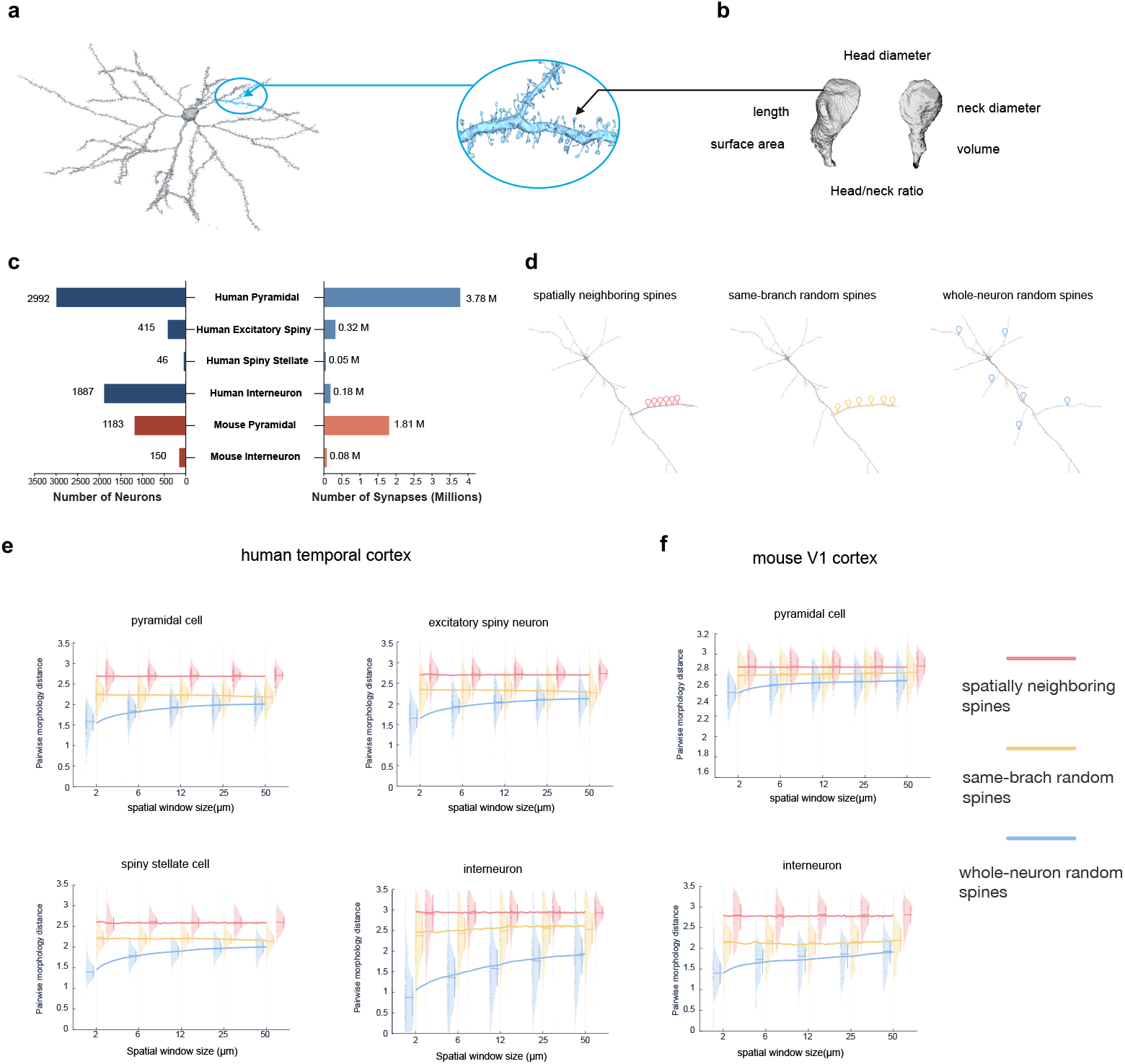
Connectomic reconstruction reveals local clustering of dendritic spine morphology. **a**,**b** High-resolution neuron reconstructions reveal dendrite and spine ultrastructure. **b**, Schematic of dendritic spine reconstruction and the 6 morphometric features used to define each spine’s position in morphology space. **c**, Connectomic sampling across datasets and reconstructed cell classes. Bars show reconstructed neuron counts (left) and analyzed spine-bearing synapse counts (right) for human left temporal cortex(H01 dataset) and mouse V1 cortex (MICrONS dataset). **d**, Schematic of the three sampling schemes used to quantify morphology clustering: spines within a local dendritic window, and randomly sampled spine-count-matched controls from either the same branch or the whole neuron. **e**,**f** Spatially neighboring dendritic spines show local morphological similarity across human and mouse cortical reconstructions. For each spatial window size, the average pairwise Euclidean distance in z-scored spine-morphology space was computed among all spines within the local window and compared with spine-count-matched branch-random and global-random controls. **e**, H01 human temporal cortex cell classes show consistently lower morphological distance in local spatial windows than in branch-random or global-random samples. **f**, The same local morphology compactness is observed in MICrONS mouse V1 pyramidal neurons and interneurons. Lower distance indicates greater morphology similarity.

### Synaptic weights exhibit center–surround organization despite local morphological clustering

Spine morphology defines the local structural context in which synapses are embedded, but spine-head volume is more directly related to synaptic efficacy and can therefore be used as an anatomical proxy for synaptic weight (*12, 22–24*). We next asked whether synaptic weights followed the same local organization as spine morphology, or whether they obeyed a distinct spatial rule. Across reconstructed H01 cell classes, synapse weight was strongly right-skewed and approximately follows lognormal distribution (Fig. 2a), consistent with prior measurements and synaptic-memory models of skewed synaptic strength distributions (*11, 23, 25*). This weight polarization was quantified using Lorenz curves and Gini coefficients across reconstructed cell classes (Gini = 0.788–0.870; Fig. 2a,c), indicating that a minority of large synapse contributed a disproportionate fraction of total synapse weight. We therefore defined large-synapse centers as the upper quintile of the spine-head-volume distribution, capturing the high-weight tail with a data-driven percentile threshold. For each large-synapse center, we measured neighboring synapse weight as a function of dendritic path distance, excluding the center synapse itself. As a branch-aware null model, we shuffled spine-head volumes within each biological branch while preserving all spine positions, branch geometry, and local spine density. Across the four analyzed H01 cell classes, neighbor weight showed a Mexican-hat-like profile around large-spine centers (Fig. 2e) and the same profile was also observed across cortical layers 2–6 (Fig. 2f). The profile contained three spatial regimes: an immediate center-proximal elevation within approximately 0–2 µm, a broad suppression annulus across intermediate distances of approximately 2–14 µm in which observed neighbor weight fell below the within-branch shuffled baseline, and a recovery or rebound zone at approximately 14–17 µm before returning toward baseline at larger distances. Thus, the spatial organization of synaptic weights around large synapses was nonmonotonic, with local elevation followed by intermediate surround suppression.

**Figure 2:**
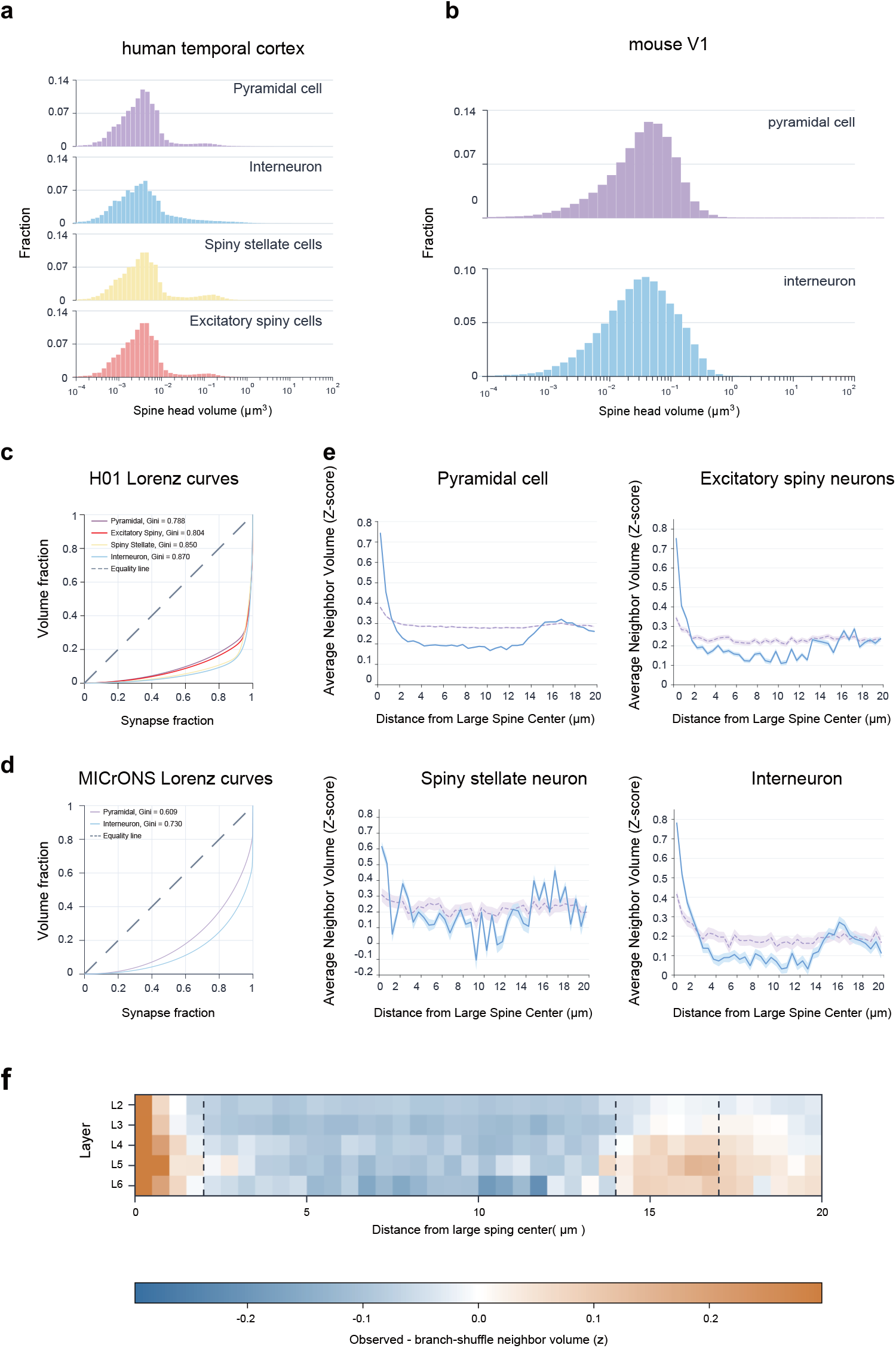
synaptic weights exhibit center-elevation/surround-inhibition organization in human temporal cortex. **a**,**b**, Synaptic weight (proxied by Spine head volume) distributions on a log scale for human temporal cortex (**a**) and mouse V1 (**b**) across reconstructed cell classes. Distributions are approximately lognormal or mixture lognormal, consistent with the known variability in synaptic size. **c**,**d**, Lorenz curves and Gini coefficients quantify polarization of synaptic weight distributions in human temporal cortex (**c**) and mouse V1 (**d**). Human temporal cortex shows stronger polarization across reconstructed cell classes than mouse V1. **e**, Center-elevation/surround-inhibition organization around large-synapse centers in human temporal cortex. Large-synapse centers were defined from the top 20% of the synaptic weight distribution, and neighboring synaptic weight was measured as a function of dendritic path distance. Across human temporal cortex cell classes, neighbor volume shows a Mexican-hat-like profile: elevation at short distances, suppression at intermediate distances, and recovery at larger distances. **f**, Heat map showing the distance-dependent neighbor synaptic weight profile across cortical layers, revealing a consistent Mexican-hat-like pattern. blue indicates suppression below the branch-shuffle baseline and orange indicates elevation.

Spatially neighboring spines shared local morphological microenvironments, whereas large synapses showed distance-dependent elevation and depletion of neighbors across a surrounding dendritic zone. Thus, synaptic weight states follow a spatial organization that cannot be inferred from local spine morphology alone. Interestingly, this spatial scale aligns with the biophysical boundaries of local dendritic microdomains, specifically the lateral spread of calcium transients along the dendritic shaft (*26, 27*). In contrast, the MICrONS mouse V1 reconstruction showed weaker spine-volume polarization (Fig. 2b,d) and did not yield a comparably robust Mexican-hat profile under the same branch-shuffle analysis (Fig. S1). The H01 result therefore points to a pronounced spatial weight-organization signature in human temporal cortex, with the MICrONS contrast suggesting that this signature varies across cortical areas and species. We propose this divergence reflects computational demands: mouse V1 primarily encodes low-dimensional, orthogonal features where functional interference is inherently minimal, whereas human temporal cortex processes high-order, highly overlapping associative representations that necessitate spatial anti-crowding to prevent catastrophic interference (*28, 29*). Furthermore, whereas structural plasticity in rodent V1 largely crystallizes following early critical periods, human temporal cortex shall support continuous lifelong learning (*14, 30*). In this context, the center–surround topology likely serves as a spatial constraint that prevents the local crowding of high-weight states, accommodating both the dense representational overlap and the sustained structural complexity of higher-order cortices.

### Spatial Hebbian learning limits co-occupation

The center–surround weight field shown in Fig. 2e suggests a spatial dimension of synaptic learning that is not captured by classical Hebbian rules or subsequent normalization-based extensions such as Oja’s rule (*1,5–8*). These models specify how correlated pre and postsynaptic activity strengthens individual synapses over time and how weight growth can be stabilized, but they do not specify how potentiated synapses are arranged along a dendritic branch. We therefore asked what spatial interaction could generate the observed short-range elevation, intermediate suppressive annulus, and long-range recovery. To formalize this topology, we added a distance-dependent spatial interaction term to a Hebbian weight dynamic.

We modeled the temporal evolution of synaptic weight *w*_*i*_ at dendritic position *i* as the sum of an activity-dependent Hebbian drive and a local spatial interaction,

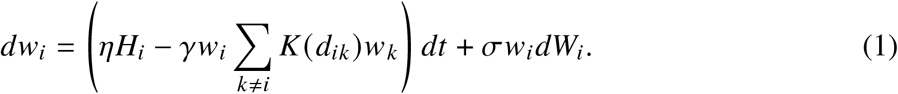

where *H*_*i*_ denotes the Hebbian drive, *d*_*ik*_ is the dendritic path distance between synapses *i* and *k, K* (*d*) is a signed spatial kernel, *γ* controls the strength of spatial coupling, and *σ* sets the multiplicative noise scale. The signed kernel captures the geometry of the observed Mexican-hat profile: a short-range attractive regime permits center-proximal co-potentiation, an intermediate inhibitory annulus penalizes nearby high-weight co-occupation, and the interaction decays toward baseline at larger distances.

Under a reduced stochastic approximation, it can be shown that the inhibitory annulus imposes a nonlinear constraint on nearby high-weight states. Specifically, if one synapse reaches a high-weight state *w*_*i*_ = *W*, the conditional probability that a neighboring synapse within the inhibitory annulus also exceeds a high-weight threshold *w*_*c*_ follows

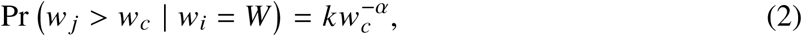

where *α* increases with the center-synapse strength *W* and depends on the spatial coupling strength, the inhibitory component of the kernel, and the noise scale; *k* is set by the lower weight bound. Thus, the spatial term does not impose a hard upper bound on individual synaptic growth. Instead, it creates a soft probabilistic ceiling for local high-weight co-occupation: weak and medium synapses can remain nearby, but the probability of another strong synapse in the inhibitory annulus falls nonlinearly as the center synapse grows (proof see methods).

We next asked whether the suppression zone reflected a reduced probability of high-weight neighbors, as predicted by the model, rather than a nonspecific change in local spine composition. To test this, we estimated the conditional occurrence probability of neighboring spines from each spine-head-volume quintile as a function of dendritic path distance from large-synapse centers (Fig. 3b,c). In the suppression annulus, the probability of upper-quintile neighbors fell below the nominal 20% expectation, reaching 18.3% in pyramidal neurons and 18.9% in spiny stellate cells; both decreases were significant under branch-preserving permutation tests (p = 0.024 and p = 0.041, respectively). Excitatory spiny cells showed the same direction of change, with upper-quintile neighbor probability reduced to 19.1%, although this decrease did not reach significance (p = 0.13). When high-weight neighbors were defined more stringently as the top decile of the synaptic weight distribution, the depletion became stronger: neighbor probability fell below the nominal 10% expectation to 8.2% in pyramidal neurons, 8.1% in excitatory spiny cells, and 9.0% in spiny stellate cells, all reaching significance (p = 0.024, 0.032, and 0.041, respectively; Fig. 3d,e). Interneurons did not show a significant effect under either threshold, consistent with their lower spine density, smaller neighbor samples per shuffle, broader permutation nulls, and reduced statistical power. Together, these results show a selective depletion of high-weight neighbors in the suppression zone across multiple cell classes, with a stronger effect under the stricter top-decile threshold, arguing against a nonspecific sampling artifact.

**Figure 3:**
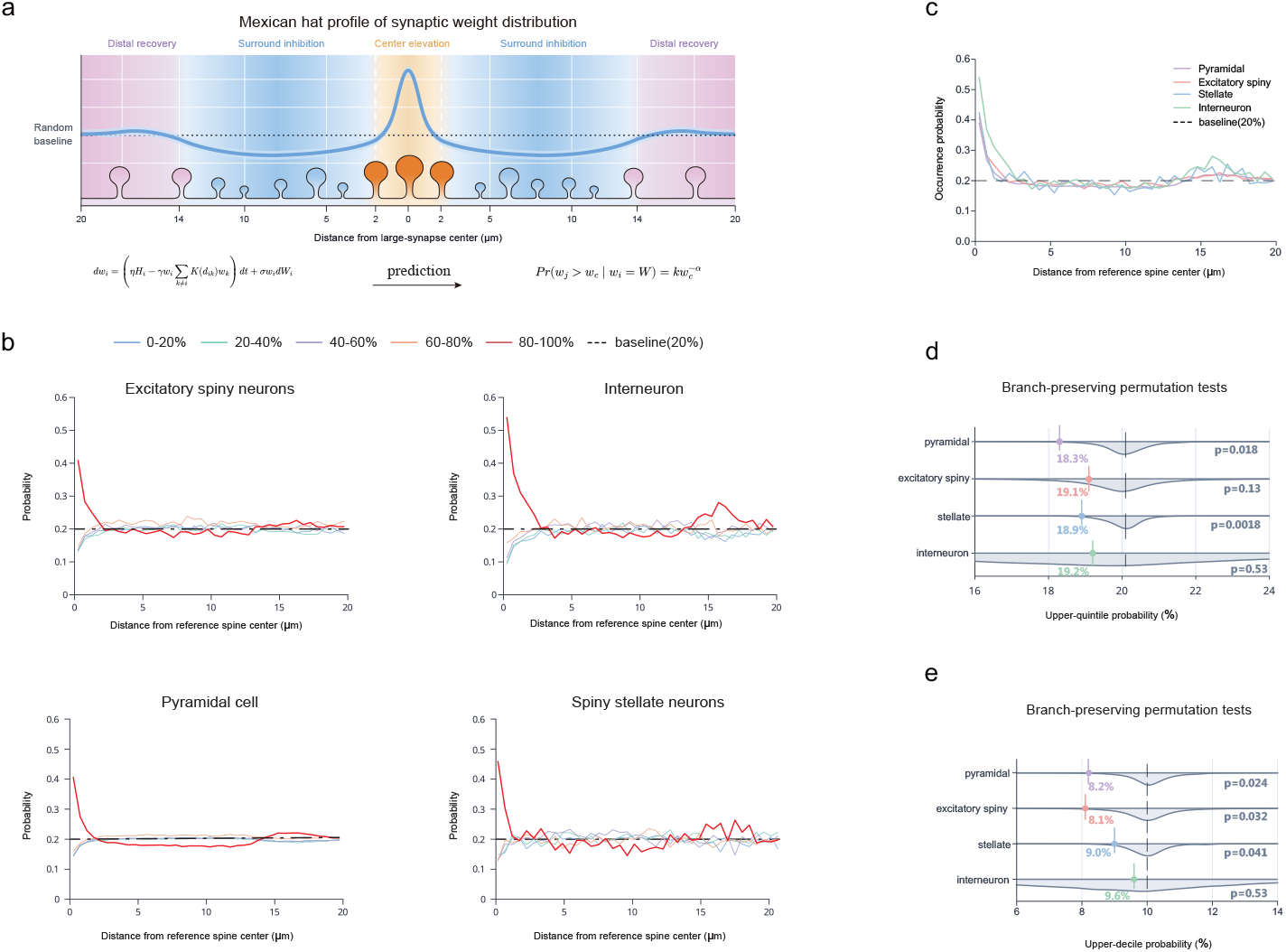
Spatially constrained Hebbian learning predicts selective high-weight suppression. **a**, Spatially constrained Hebbian model. A Hebbian drive promotes activity-dependent synaptic growth, whereas a distance-dependent interaction term imposes spatial motif. In the inhibitory surround, the model predicts a nonlinear reduction in the probability that a neighboring synapse also enters a high-weight state. **b**, Neighbor composition around top 20% large synapse centers across four H01 cell classes. Neighboring synapses were grouped by synaptic weight quintile and plotted as a function of dendritic path distance. The dashed line marks the nominal 20% fraction for each quintile. **c**, Distance-dependent upper-quintile neighbor frequency around large-synapse centers across human temporal cortex cell classes. **d**, Permutation test for selective high-weight suppression in the suppression zone. Spine head volumes (proxy for synaptic weight) were randomly reassigned among spines within each branch, while spine positions were held fixed. Observed large neighbor (top 20%) probabilities around large-synapse centers were compared with 1,000 branch-preserving shuffles. **e**, Same branch-preserving permutation analysis as in **d**, but with large-synapse neighbors defined as the top 10% rather than the top 20% of the synaptic weight distribution.

This probability analysis indicates that the surround component of the Mexican-hat profile does not reflect a nonspecific reduction of nearby synapses, but rather a selective decrease in the probability that nearby synapses enter high-weight states. Thus, synaptic weight appears not to be governed by temporal Hebbian drive alone; instead, the transition into and maintenance of high-weight states are also constrained by distance-dependent spatial interactions.

### Spatial regularization reduces forgetting by reorganizing learned-direction geometry

In biological dendrites, physical proximity can reflect functional proximity: nearby spines are more likely to receive synchronized or feature-related inputs from upstream neurons, forming local dendritic input clusters (*31–33*). This relationship motivates an abstraction from dendritic distance to distance among learned directions. We therefore translated the center-elevation/surround-suppression synaptic weight profile into Spatial Synaptic Regularization (SSR), a regularizer for artificial networks. SSR tests whether a center–surround competition prior can reduce representational crowding by discouraging multiple high-amplitude learned directions from occupying the same local neighborhood in representation space.

We first tested SSR in replay-free task-incremental image classification with a ResNet-18 backbone (*34*). Across Split-CIFAR-100, Split-CIFAR-10, Split-TinyImageNet, and a five-dataset cross-domain stream, tuned SSR paired with knowledge distillation (SSR+) improved the accuracy– forgetting trade-off relative to the strongest matched comparator evaluated under each continual-classification protocol (Fig. 4). On Split-CIFAR-100, the accuracy-selected SSR+ run reached 73.50±0.04% average accuracy and 3.05±0.04 average forgetting, exceeding LwF (72.44±0.18%, 8.14±0.25) and matched same-protocol geometry controls in average accuracy. In a separate retention-control gate, SSR+ reached 72.68±0.14% and 1.48±0.09, giving the lowest mean forgetting when SSR+ was compared with KD-only, cosine orthogonality, center loss, supervised contrastive loss, and other same-code controls under the same seed set and budget (Table S1). Thus, Fig. 4a,b report matched trade-off points and positive accuracy–forgetting margins across all four streams, while the supplementary control gate verifies that the same SSR+ implementation also gives the strongest forgetting reduction under matched C100 settings.

**Figure 4:**
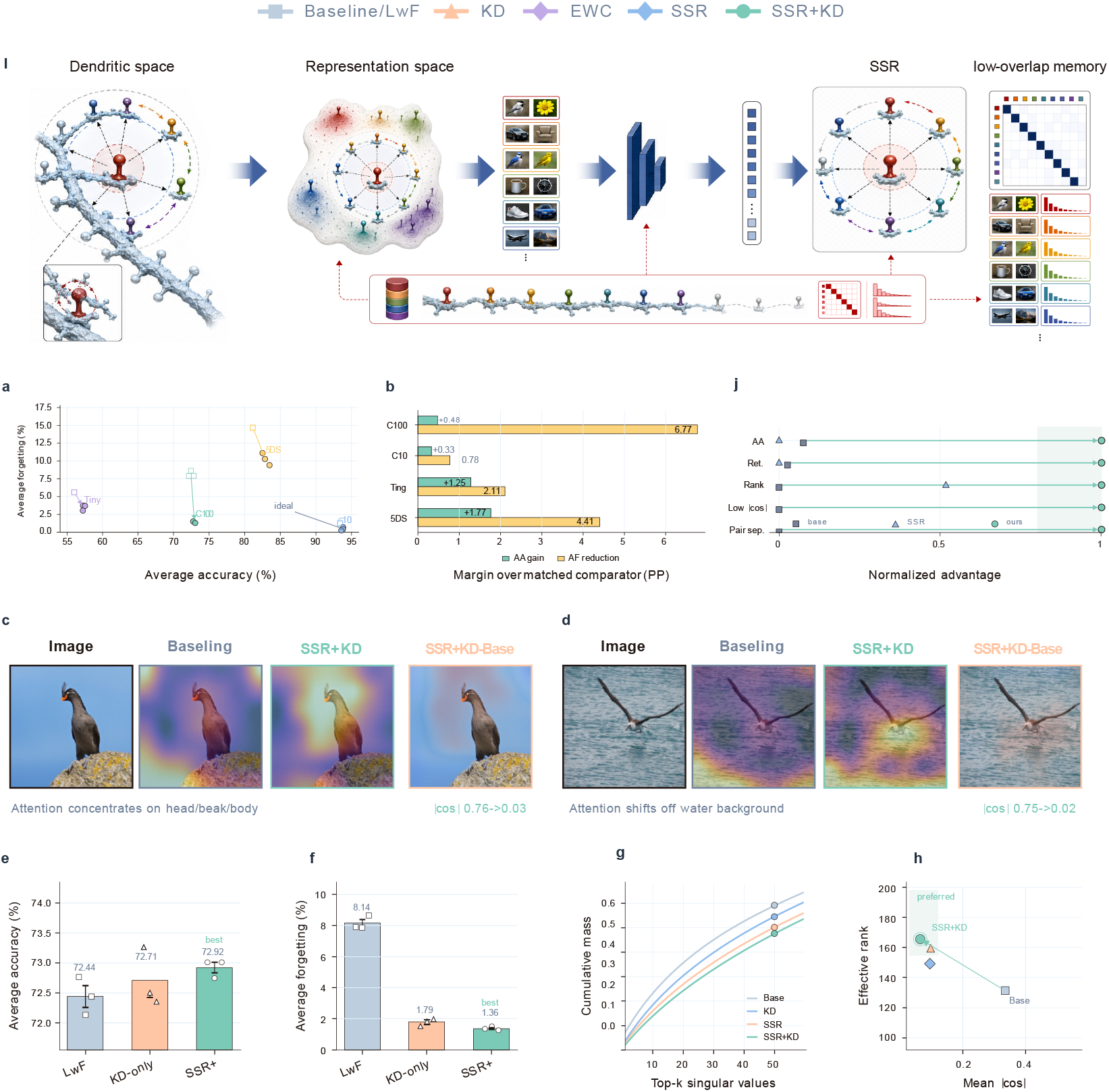
SSR improves continual-classification retention and yields high-rank low-overlap memories. **a**, SSR architecture from task-stream features to low-overlap memory. **b**, Multi-metric dominance profile for matched CUB gains. **c**,**d**, Matched trade-off shifts and margins across four task streams. **e**,**f**, CUB CAM examples show stronger bird-region localization and lower hard-pair prototype cosine. **g**,**h**, Seed-level C100 average accuracy and forgetting use the matched SSR+ run and matched baselines. **i**, Slower CUB singular-value mass accumulation indicates a less compressed, higher-rank classifier geometry. **j**, Rank–overlap map summarizes the target state, where higher rank and lower overlap are preferred.

**Figure 5:**
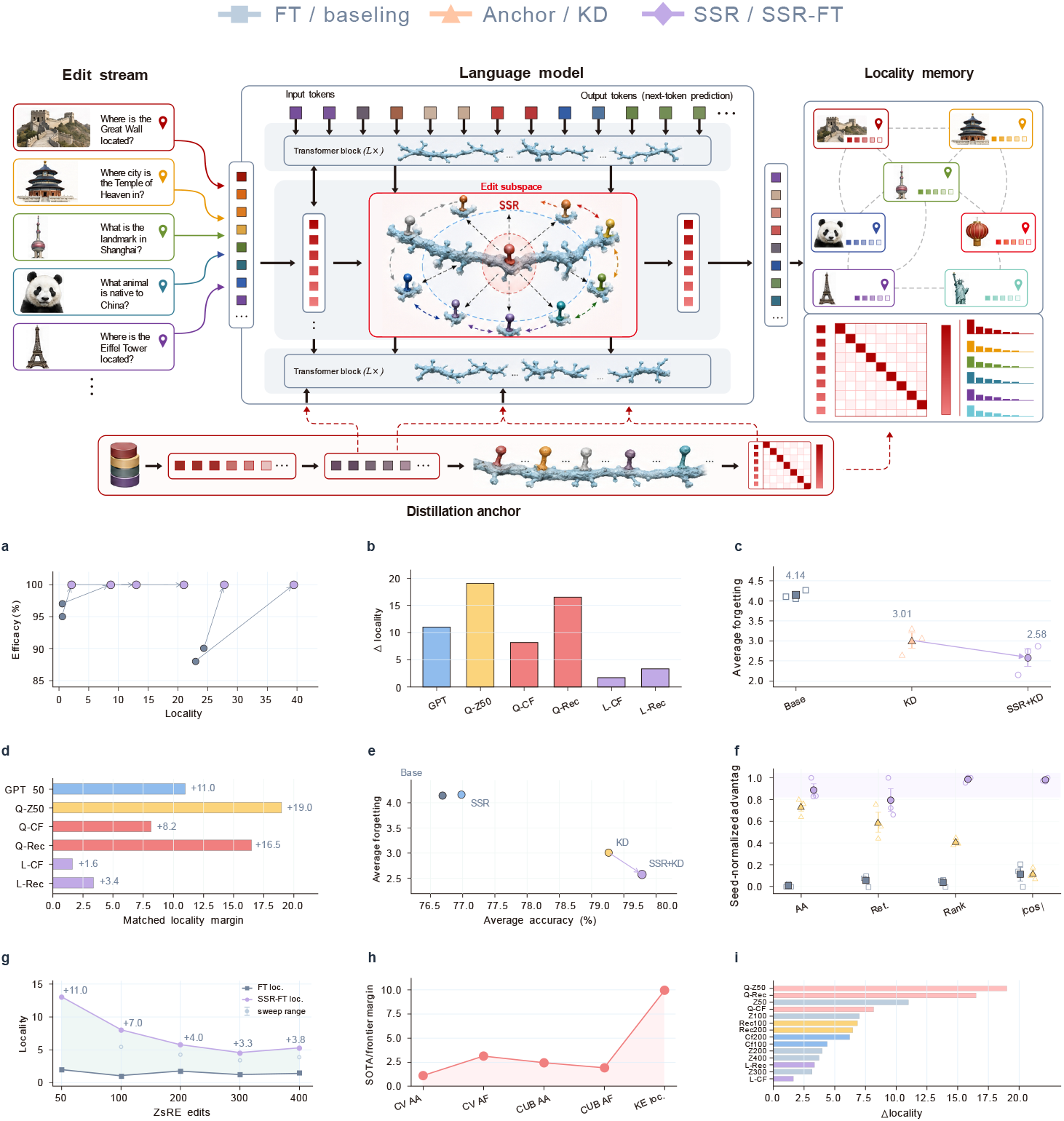
SSR transfers to matched editing and adapter probes. **a**, SSR edit-stream architecture. **b**, Locality–efficacy scatter for matched edits. **c**, Locality gains across GPT, Qwen2.5, and Llama slices. **d**, Seed-level adapter forgetting. **e**, FT-style editing margins. **f**, Adapter accuracy–forgetting trade-off. **g**, Adapter geometry profile. **h**, ZsRE edit-budget scaling. **i**, Cross-domain matched-protocol summary. **j**, Knowledge-editing gain atlas.

The same matched trade-off pattern persisted in the other streams. On Split-CIFAR-10, SSR+ reached 93.72±0.07% average accuracy and 0.45±0.13 forgetting, compared with 93.39±0.18% and 1.23±0.25 for LwF. On Split-TinyImageNet, SSR+ reached 57.31±0.10% average accuracy and 3.47±0.23 forgetting, compared with 56.03±0.37% and 5.59±0.33 for LwF. In the five-dataset benchmark, SSR+ reached 82.92±0.28% average accuracy and 10.28±0.49 forgetting, exceeding ER-ACE (81.15±0.12% and 14.68±0.35). Thus, adding the spatial competition prior to distillation improves the matched accuracy–forgetting trade-off across the evaluated task-incremental streams. Cached run summaries for the selected SSR+ configurations provide an initial training-cost scope (Table S2), but these logs were collected from heterogeneous cluster runs and are not treated as a hardware-normalized speed benchmark.

### SSR yields high-rank, low-overlap memory geometry

We next asked what representational change accompanied the retention gain. On CUB-200-2011 (*35*), SSR+ gave the best matched 20-task fine-grained classification trade-off among the evaluated methods, improving average accuracy from 78.68% to 81.10% and reducing forgetting from 3.49 to 1.58. The same run increased classifier effective rank from approximately 131 to 166 and reduced mean off-diagonal prototype cosine from 0.335 to 0.064. These diagnostics indicate that SSR+ preserves a higher-rank, lower-overlap class-memory geometry rather than merely shrinking updates. Across the matched ablation, SSR supplied the structural anti-crowding prior and distillation supplied the functional anchor, with the combined SSR+ configuration giving the strongest retention point.

Additional visual and dynamical diagnostics identify the mechanism behind this matched trade-off. Attention maps from fine-grained CUB examples show that SSR+ concentrates more spatial evidence on diagnostic bird regions such as the head and beak, consistent with a less diffuse class representation. At task resolution, the same SSR+ run gave the lowest mean update norm and the most positive lag-1 update cosine among the matched CUB variants. We therefore use rank, prototype overlap, attention localization, and task-level temporal persistence as the main interpretability evidence, while retaining matrix-internal Fourier spectra as a supplementary scope analysis in Fig. S3.

### The spatial constraint generalizes beyond classifiers to language-model editing and adaptation

Finally, we used two controlled transfer probes to ask whether the same anti-crowding prior can stabilize learned directions outside classifier-head continual learning. First, in FT-style KnowEdit streams, SSR-FT improved the locality–efficacy trade-off relative to true fine-tuning within the evaluated editor family. On GPT-2 XL ZsRE, SSR-FT improved locality from 2.0 to 13.0 while retaining ceiling 100% efficacy in the 50-edit setting and maintained positive locality margins in longer 100–400 edit streams. In the completed Qwen2.5-VL-3B and Llama-3-8B editing pairs reported in Table 1, SSR-FT retained ceiling efficacy or improved efficacy while also increasing locality. The largest open-checkpoint gains appeared on Qwen2.5-VL-3B, where ZsRE-50 locality increased from 2.0 to 21.0 at ceiling efficacy and WikiRecent-100 locality increased from 22.96 to 39.46 while efficacy improved from 88.0% to 100.0%. These editing results should be interpreted as matched fine-tuning-style probes, not as a broad comparison with specialized model-editing systems. Because large-language-model attention maps are not directly comparable to class-localized visual evidence, the LLM evidence is reported through locality–plasticity trade-offs and update-subspace diagnostics rather than through attention visualization.

**Table 1:** Matched-protocol summary for SSR across classification, geometry, knowledge editing, and adapters. Each row reports the matched comparator → the SSR configuration within the same protocol. Margins are expressed in the preferred direction; red., reduction. AA, average accuracy; AF, average forgetting; Eff., efficacy; Loc., locality. LLM editing and adapter rows are controlled transfer probes rather than exhaustive literature benchmarks.

| Setting | Pair | Primary readout | Margin | Secondary readout | Margin |
| --- | --- | --- | --- | --- | --- |
| Split-CIFAR-100 | LwF → SSR+ | AA ↑ 72.44±0.18 → **73.50±0.04** | +1.06 | AF ↓ 8.14±0.25 → **3.05±0.04** | 5.09 red. |
| Split-CIFAR-10 | LwF → SSR+ | AA ↑ 93.39±0.18 → **93.72±0.07** | +0.33 | AF ↓ 1.23±0.25 → **0.45±0.13** | 0.78 red. |
| Split-TinyImageNet | LwF → SSR+ | AA ↑ 56.03±0.37 → **57.31±0.10** | +1.28 | AF ↓ 5.59±0.33 → **3.47±0.23** | 2.11 red. |
| 5-dataset stream | ER-ACE → SSR+ | AA ↑ 81.15±0.12 → **82.92±0.28** | +1.77 | AF ↓ 14.68±0.35 → **10.28±0.49** | 4.41 red. |
| CUB200 classification | Base → SSR+KD | AA ↑ 78.68 → **81.10** | +2.42 | AF ↓ 3.49 → **1.58** | 1.90 red. |
| CUB200 geometry | Base → SSR+KD | Eff. rank ↑ 131.37 → **165.69** | +34.32 | \|cos\| ↓ 0.335 → **0.064** | 0.271 red. |
| GPT-2 XL ZsRE-50 | FT → SSR-FT | Eff. ↑ **100.0** → **100.0** | ceiling | Loc. ↑ 2.0 → **13.0** | +11.0 |
| Qwen2.5-VL-3B ZsRE-50 | FT → SSR-FT | Eff. ↑ **100.0** → **100.0** | ceiling | Loc. ↑ 2.0 → **21.0** | +19.0 |
| Qwen2.5-VL-3B CF-100 | FT → SSR-FT | Eff. ↑ 97.0 → **100.0** | +3.0 | Loc. ↑ 0.51 → **8.69** | +8.18 |
| Qwen2.5-VL-3B Recent-100 | FT → SSR-FT | Eff. ↑ 88.0 → **100.0** | +12.0 | Loc. ↑ 22.96 → **39.46** | +16.50 |
| Llama-3-8B CF-100 | FT → SSR-FT | Eff. ↑ 95.0 → **100.0** | +5.0 | Loc. ↑ 0.51 → **2.15** | +1.64 |
| Llama-3-8B Recent-100 | FT → SSR-FT | Eff. ↑ 90.0 → **100.0** | +10.0 | Loc. ↑ 24.41 → **27.77** | +3.36 |
| Low-rank adapter | KD → SSR+KD | AA ↑ 79.26±0.16 → **79.78±0.20** | +0.52 | AF ↓ 3.01±0.19 → **2.58±0.22** | 0.43 red. |

Second, we implemented a low-rank adapter experiment as a parameter-efficient adaptation probe (*36*). Using frozen CUB200 ResNet-18 features, we trained a rank-16 bottleneck adapter and classifier head over the same 20-task stream. SSR was applied both to class prototypes and to adapter output-basis directions. Across three seeds, SSR+KD improved the matched adapter trade-off relative to KD-only, increasing average accuracy from 79.26 to 79.78 and reducing forgetting from 3.01 to 2.58, while increasing effective rank from 160.15 to 168.08 and reducing prototype cosine from 0.101 to 0.075. This result supports SSR as a transferable geometry prior within the evaluated frozen-feature adapter protocol, but it should not be read as a general LoRA benchmark.

## Discussion

Our results suggest that synaptic learning is not only a temporal process but also a spatially organized one. Classical Hebbian rules and their extensions specify when synapses change and how strongly they grow, treating each synapse as an isolated variable. The center-elevation/surround-inhibition topology identified here adds a complementary principle: a synapse entering a high-weight state alters the local probability landscape for its neighbors, permitting co-potentiation in its immediate vicinity, while constraining high-weight co-occupation across the intermediate surrounding suppression zone. Critically, this constraint was selective rather than uniform. Low and medium weight neighbors were weakly affected, while large neighbors were inhibited. This indicates that neurons do not simply suppress local plasticity but spatially organize synaptic states, offering a route to the stability-plasticity dilemma in which stability arises from regulating allocation of weights.

Although static electron microscopy cannot directly observe synaptic plasticity as it unfolds, large scale connectomic structure can reveal the statistical footprint of learning rules. Statistical physics offers a useful analogy: effective interaction potentials are often inferred from equilibrium pair correlations in static particle configurations, rather than from direct observation of individual particle trajectories. The spatial scale of the suppression zone, spanning approximately 10–20 µm, aligns with known boundaries of lateral Ca^2+^ diffusion and dendritic integration (*11, 13, 15, 31, 37*), raising the possibility that local biophysical compartmentalization structurally implements the spatial organization we observe statistically. Definitive confirmation of the underlying mechanism and the spatially constrained Hebbian rule will require dynamic experiments, such as in vivo two-photon imaging of synaptic weight changes alongside localized perturbation of Ca^2+^ signaling.

SSR algorithm extends this center–surround principle to representation space, permitting co-occupation of similar learned directions while penalizing high-amplitude overlap at intermediate distances. Unlike classical regularization, replay, or distillation — which protect specific parameters, examples, or functions (*38–43*) — SSR constrains the relational geometry among learned states themselves. Because this constraint acts on geometry rather than on any task-specific output, it could be applied to classifier prototypes, editing subspaces, and adapter bases, all of which share a common failure mode in which new high-amplitude directions overlap with existing ones. Forgetting in connectionist networks is often attributed to insufficient representational capacity or to the absence of protected parameters, motivating solutions that expand capacity or freeze selected weights. Our results point to a complementary diagnosis: interference can arise even when capacity is sufficient, as high amplitude directions are allowed to occupy the same neighborhood without spatial constraint. Stability, in this view, depends not only on how much representational space exists, but on how it is organized. This allows stability to be achieved without freezing plasticity, converging with the biological topology observed in human cortex.

Several limitations remain. The center-surround topology was absent in the analyzed MICrONS mouse V1 reconstruction, and additional connectomes spanning more cortical areas, species, and developmental stages will be needed to determine its generality. On the artificial side, the present experiments provide matched performance, geometry, and preliminary run-cost summaries, but they do not yet constitute a hardware-normalized efficiency study or a broad benchmark across all continual-learning and model-editing families. These limitations do not change the central implication: learning systems may achieve stability by organizing the spatial distribution of synaptic weight states, rather than by restricting plasticity itself.

## Funding

This work was supported in part by Shanghai Gaofeng team support. Full grant identifiers will be provided by the authors at submission.

## Author contributions

H.Z. and Y.C. contributed equally to this work. H.Z. and Y.C. designed the artificial-network experiments, generated figures, and prepared the visualization workflows. All authors interpreted the results and wrote the manuscript.

## Competing interests

The authors declare no competing interests.

## Data, code and materials availability

Code for reproducing the artificial-network experiments, including training entry points, selected configuration files, metric aggregation scripts, LLM-editing probes, and AI figure workflows, is available at https://github.com/ydchen0806/ssr-ai-reproducibility. The repository documents the public AI datasets used in the experiments and separates generated results, checkpoints, and downloaded data from version-controlled source code.

## Supplementary materials

Materials and methods

Appendix S1

Figures S1 to S3

Tables S1 to S7

## Materials and Methods

### Connectomics datasets and synapse-level measurements

The biological analysis used the H01 human temporal cortex reconstruction and the MICrONS mouse visual-cortex reconstruction (*17, 19–21*). Synapses were assigned to dendritic arbors when reconstructed synaptic contacts, postsynaptic compartments, and dendritic path distances were available. For each synapse *i*, the analysis stored a dendritic coordinate *s*_*i*_, a spine morphological feature vector *m*_*i*_, and a spine-head-volume proxy *v*_*i*_ for synaptic weight. Spine volume is an indirect measure of synaptic efficacy, but its correlation with postsynaptic density and synaptic morphology makes it a standard anatomical proxy for large-scale EM analysis (*12, 22–24*).

For a pair of synapses on the same dendritic branch, dendritic distance was computed as *d*_*i j*_ = |*s*_*i*_ − *s* _*j*_ | along the branch path rather than as Euclidean distance. Local morphology similarity was summarized by

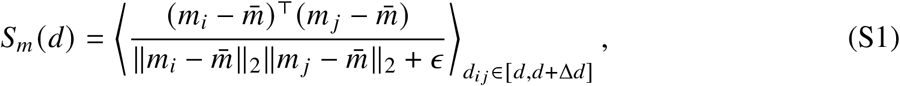

where brackets denote an average over synapse pairs in a distance bin. Weight organization was measured by the corresponding binned covariance or correlation of log-transformed spine volumes,

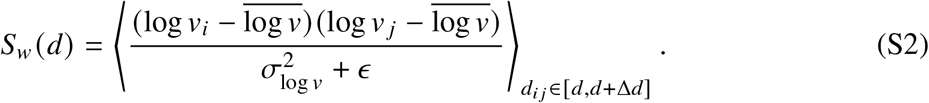

The qualitative claim in the main text is the sign dissociation between *S*_*m*_ (*d*) and *S*_*w*_ (*d*) in H01: neighboring spines are morphologically related, but nearby large-volume synapses are not simply co-clustered.

**Figure S1:**
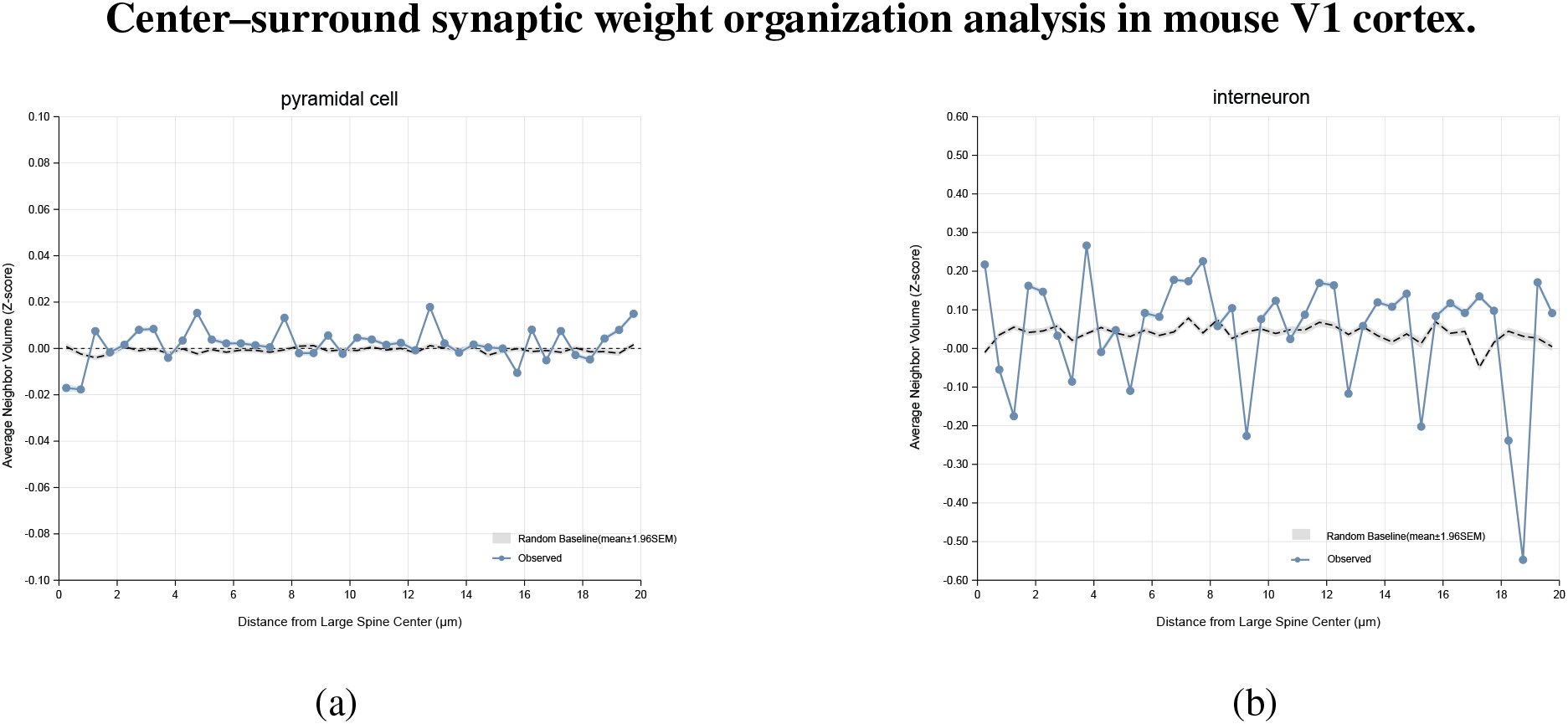
Average neighbor synaptic weight as a function of dendritic path distance from large-synapse centers (top 20% volume quintile) in mouse V1. Analyzed under the identical pipeline used for human H01 data (Fig. 2e), the mouse V1 data does not exhibit a robust center–surround topology, remaining largely flat relative to the branch-shuffled baseline across the 0–20 µm range.

### Spatially regularized Hebbian model

The derivation below analyzes a reduced stochastic Hebbian system with distance-dependent interaction and multiplicative noise. It is used as a mechanistic reduction showing how a high-weight synapse can lower the probability of nearby high-weight co-occupation under explicit mean-field, frozen-strong-synapse, strong-inhibition, and reflecting-floor assumptions.

We consider *N* synapses on a dendrite whose weights are updated by Hebbian learning under spatial interactions. The temporal evolution of the weight *w*_*i*_ > 0 of the *i*-th synapse is jointly driven by deterministic drift forces, including Hebbian drive and spatial interactions, and multiplicative Gaussian white noise, representing molecular and receptor-level thermal fluctuations. The dynamics are described by

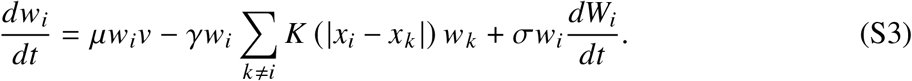

Equivalently,

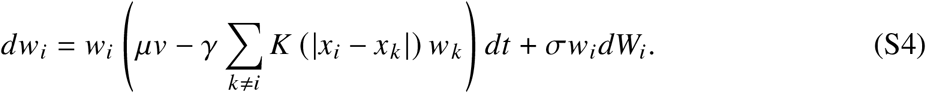

Here, 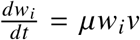 denotes the classical Hebbian learning process, µ is the Hebbian learning rate, and *v* denotes the postsynaptic neuronal activity. The term

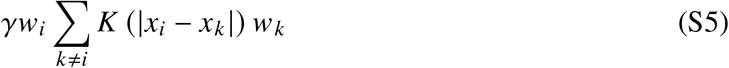

is the spatial regularization term describing interactions between synapses. The function *K* (*d*) is the spatial coupling function, *γ* denotes the global spatial coupling strength, *σw*_*i*_ *dW*_*i*_ is the multiplicative Gaussian noise term, *σ* is the noise intensity, and *W*_*i*_ denotes an independent standard Wiener process.

We next consider a pair of synapses *i* and *j* whose spatial distance lies within the inhibitory interval, that is,

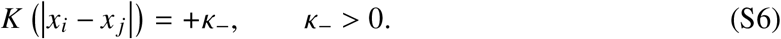

Using a mean-field approximation, the total interaction contributed by the remaining *N* −2 synapses is treated as a homogeneous background inhibitory field,

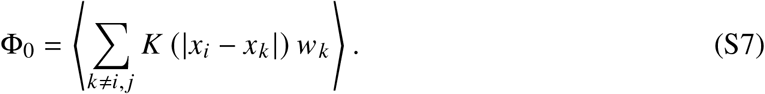

The original high-dimensional system is therefore reduced to a two-dimensional nonlinear coupled stochastic differential equation (SDE):

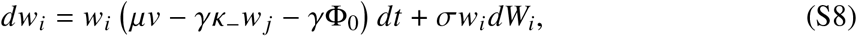

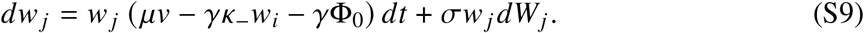

Because the original equation contains a multiplicative noise term, *σwdW*, we introduce the coordinate transformation

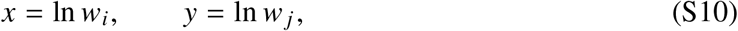

to decouple the fluctuation term and facilitate subsequent analysis.

By Itô’s lemma, for *f* (*w*) = ln *w*, its differential is

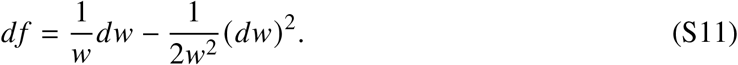

Substituting the expression for *dw*_*i*_ and applying the quadratic variation rule (*dW*_*i*_)^2^ = *dt*, the system is transformed into

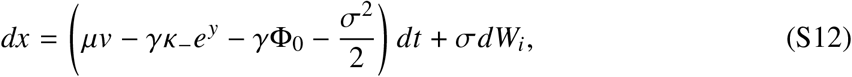

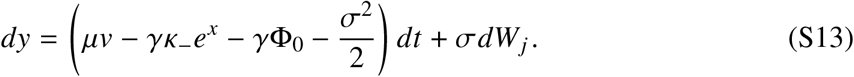

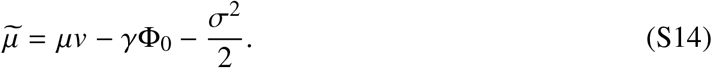

We then obtain the additive-noise system

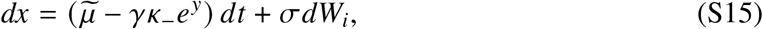

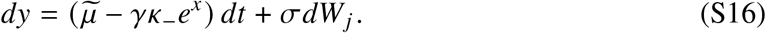

We next examine the effect of a strong synapse on its neighbor. Suppose that synapse *i* reaches an extremely large weight, so that

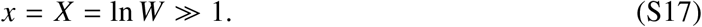

Because *e*^*X*^ is a dominant term, the deterministic drift of *y* is strongly governed by negative feedback:

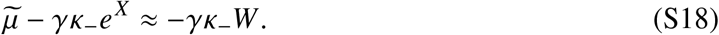

This implies that the characteristic relaxation time of *y* satisfies

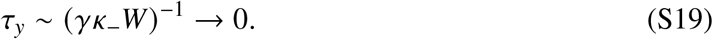

Therefore, *y* rapidly relaxes to a small state, *e*^*y*^ → 0, causing the coupling term *γκ*_−_*e*^*y*^ in the dynamics of *x* to vanish. The slow variable *x* can then be adiabatically eliminated and treated as a frozen parameter.

Because large and small synapses evolve on separated time scales, the system mathematically forms a singular perturbation problem. In this limit, the conditional dynamics of *y* reduce to Brownian motion with a constant drift:

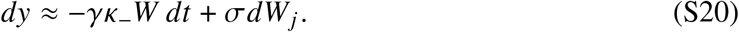

To obtain the steady-state distribution of the physical weight *w* _*j*_, we again use Itô’s formula to apply the inverse transformation *w* _*j*_ = exp(*y*). This transformation introduces a positive second-order drift correction, 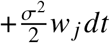, yielding the effective SDE of *w*_j_ conditioned on *W*:

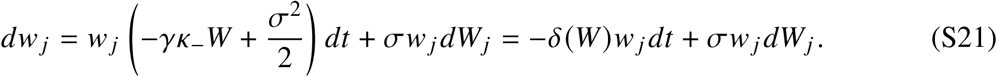

Here,

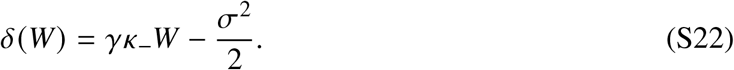

When *W* is sufficiently large, *δ*(*W*) > 0.

Because of stochasticity, it is difficult to analyze the evolution of an individual synaptic weight directly. We therefore examine the probability distribution that the weights of multiple synapses should follow. The steady-state probability density function, or invariant measure, of synapse *w* _*j*_, denoted by *p*, satisfies the Fokker–Planck equation

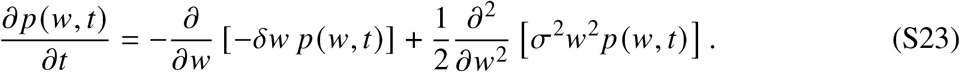

Synaptic strength has a lower bound *w*_min_, with *w* ≥ *w*_min_ ≥ 0. After a period of development, the neuron tends toward a stable state and the network approaches a steady state. At this point, the probability density no longer changes macroscopically over time. The system therefore satisfies the zero-probability-flux condition:

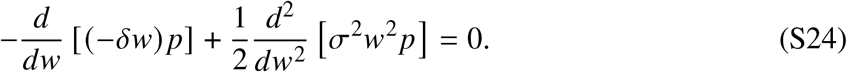

Integrating once, the equation reduces to a first-order ordinary differential equation:

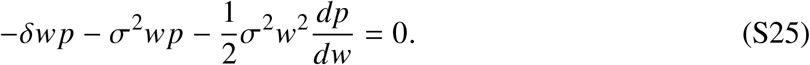

Separating variables and simplifying gives

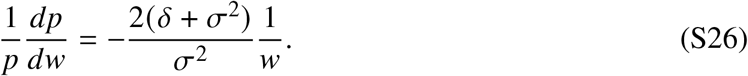

Integrating this expression shows that *w* _*j*_ follows a truncated inverse power-law distribution:

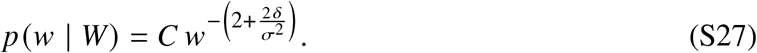

Using the normalization condition

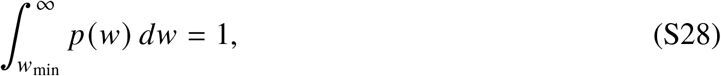

we obtain the partition constant

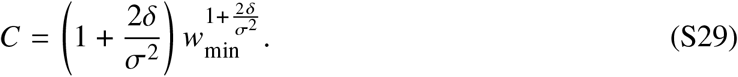

For an arbitrary threshold *w*_*c*_ > *w*_min_, the conditional tail probability, namely the probability of co-occurrence of large synapses, is

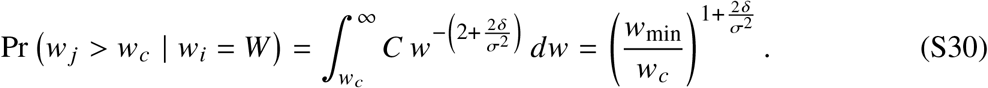

Substituting the derived effective drift coefficient

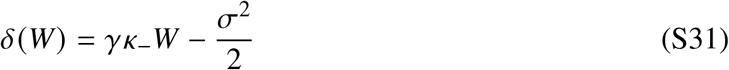

into the power-law exponent, we obtain

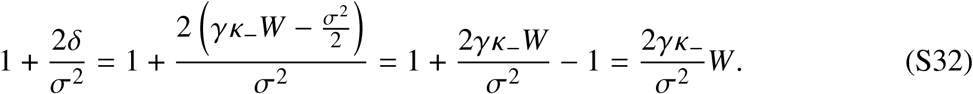

Therefore, in a synaptic dynamical system driven by Hebbian learning, with multiplicative noise and spatial interactions, if one synapse reaches an extremely large weight *w*_*i*_ = *W* ≫ 1, then the conditional probability that any synapse *w* _*j*_ within its intermediate-range neighborhood crosses a fixed threshold *w*_*c*_ obeys the following asymptotic decay law:

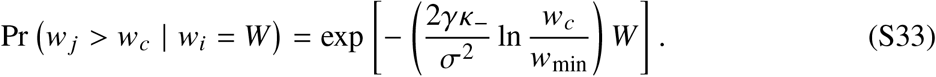

For fixed *W* and *w*_min_, this can also be written as

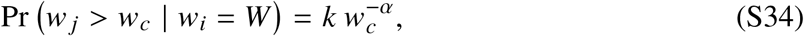

Where

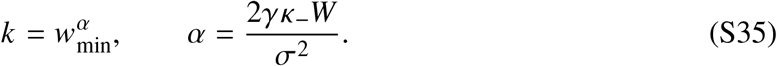

### SSR objective for artificial networks

Let *q*_*i*_ and *q* _*j*_ be two learnable directions. Depending on the experiment, *q*_*i*_ is a classifier prototype, a decoder-channel vector, a mask class embedding, an editing-subspace direction, or an adapter output-basis direction. We normalize directions before computing cosine distance,

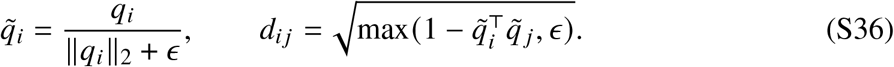

The SSR loss over a set Q of *n* directions is

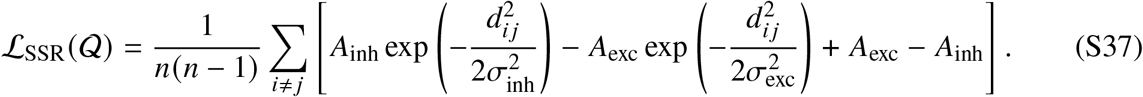

The constant shift makes the diagonal-compatible region nonnegative after removing self-pairs and does not affect gradients among distinct directions. For class-incremental streams, the loss is applied only to currently observed or previously observed class prototypes rather than to unseen classes.

SSR differs from generic orthogonality or prototype decorrelation controls because the pairwise penalty is not a monotonic push toward global separation. The difference-of-Gaussians form imposes a signed center–surround geometry: very near compatible directions are not treated the same way as directions in the inhibitory annulus, and distant directions interact weakly. The matched controls in Table S1 therefore test whether this structured annulus prior adds information beyond ordinary pairwise separation, covariance decorrelation, center compactness, or supervised contrastive geometry.

The main SSR+ objective is

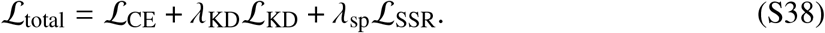

For old-task examples or current-task examples evaluated by a frozen teacher, distillation is

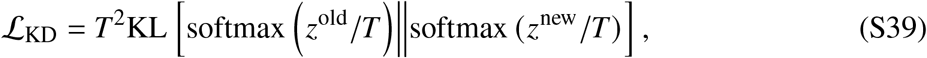

where *T* is the temperature. This decomposition is important for interpretation: KD supplies the functional anchor, whereas SSR supplies a structural anti-overlap prior (*38, 39*).

### Continual-learning protocols and metrics

For a stream of *T* tasks, let *a*_*t,k*_ be accuracy on task *k* after training task *t*. Average accuracy after the final task is

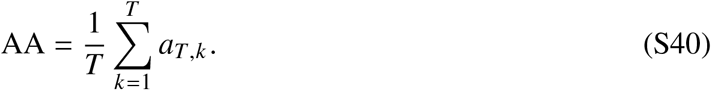

Average forgetting is

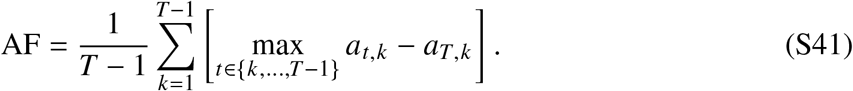

These definitions follow standard continual-learning practice (*44, 45*). Split-CIFAR-10, Split-CIFAR-100, Split-TinyImageNet, and the five-dataset stream used ResNet-18 backbones (*34,46,47*). The five-dataset stream followed the sequence CIFAR-10, MNIST, FashionMNIST, SVHN, and CIFAR-100. Classical baselines included EWC, SI, MAS, LwF/KD, and replay-style references (*39–42, 48–53*).

**Table S1:** Matched Split-CIFAR-100 geometry-control ablation. All rows use the same task stream, seed set, training budget, optimizer family, and distillation strength where applicable. Values are mean ± s.d. over three seeds. This table reports a retention-focused SSR+ configuration; the accuracy-selected SSR+ point plotted in Fig. 4a,b is reported in the main numerical matched-protocol table.

| Method | Question tested | AA ↑ | AF ↓ |
| --- | --- | --- | --- |
| LwF / KD anchor | Distillation reference under the matched task stream | 72.44±0.32 | 8.14±0.44 |
| **SSR+** | Center-surround spatial prior paired with KD | 72.68±0.24 | **1.48±0.16** |
| KD-only matched path | SSR+ training path with the spatial term disabled | 72.71±0.48 | 1.79±0.23 |
| Cosine orthogonality | Generic pairwise cosine separation | 72.44±0.22 | 1.77±0.22 |
| Prototype decorrelation | Generic covariance or decorrelation control over proto-types | 72.37±0.36 | 1.90±0.38 |
| Spectral regularization | Generic spectrum-shaping regularizer | 72.13±0.18 | 1.80±0.39 |
| Center loss | Class-center compactness control | 72.69±1.18 | 1.59±0.04 |
| Supervised contrastive loss | Standard contrastive geometry control | 72.71±0.70 | 1.77±0.40 |
| KD plus weight decay | Optimizer-level shrinkage control | 31.69±0.56 | 39.04±0.33 |

### Computational-cost scope from existing run summaries

The artificial-network experiments retained per-seed <monospace>summary.json</monospace> files containing average accuracy, average forgetting, seed, task count, epoch count, and wall-clock training time. We use these saved times only as a feasibility and cost-scope audit, because the historical jobs were executed under heterogeneous cluster loads and were not designed as a hardware-normalized speed benchmark. SSR adds no inference-time parameters in the classifier-head experiments because the SSR term is a training loss over learned directions and is removed at evaluation.

### Representation-geometry diagnostics

For a classifier matrix *W* ∈ ℝ^*C*×*D*^ with singular values *σ*_*i*_, we used the normalized spectrum *p*_*i*_ = *σ*_*i*_/∑ _*j*_ *σ*_*j*_ and effective rank

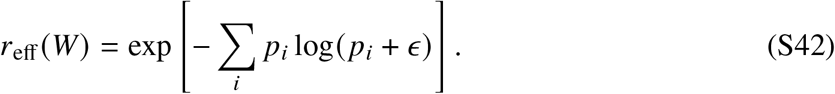

Prototype overlap was measured as the mean absolute off-diagonal cosine,

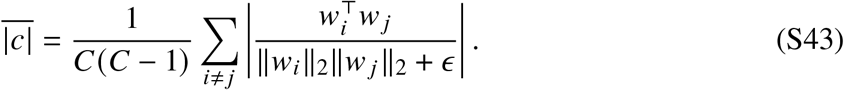

**Table S2:**
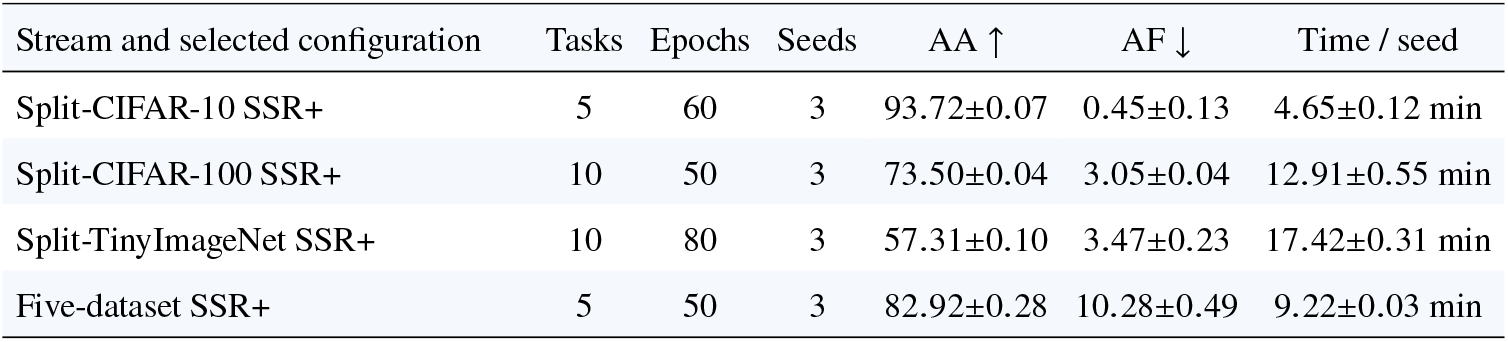
Cached run-cost scope for selected SSR+ continual-classification configurations. Values are mean ± s.e.m. over three seeds from existing training summaries. The times are saved wall-clock training times from historical cluster runs and should be interpreted as feasibility scope rather than as a hardware-normalized speed comparison. Inference overhead is none for these classifier-head experiments because the SSR term is used only during training.

| Stream and selected configuration | Tasks | Epochs | Seeds | AA ↑ | AF ↓ | Time / seed |
| --- | --- | --- | --- | --- | --- | --- |
| Split-CIFAR-10 SSR+ | 5 | 60 | 3 | 93.72±0.07 | 0.45±0.13 | 4.65±0.12 min |
| Split-CIFAR-100 SSR+ | 10 | 50 | 3 | 73.50±0.04 | 3.05±0.04 | 12.91±0.55 min |
| Split-TinyImageNet SSR+ | 10 | 80 | 3 | 57.31±0.10 | 3.47±0.23 | 17.42±0.31 min |
| Five-dataset SSR+ | 5 | 50 | 3 | 82.92±0.28 | 10.28±0.49 | 9.22±0.03 min |

The CUB200 mechanism analyses in Fig. 4 use the same quantities and include class pairs with high baseline prototype cosine that become separated under SSR+KD. The normalized structural-functional matrix combines effective rank, 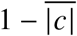, final AA, and inverse AF.

### CUB200 classification, capacity, and masks

CUB-200-2011 was used as the main fine-grained continual-learning benchmark (*35*). Classification experiments used ImageNet-pretrained ResNet-18 frozen features and a sequential classifier over 20 tasks with 10 classes per task. The semantic-order result reported in the main text is the three-seed summary: baseline 78.68 ± 0.39 AA and 3.49 ± 0.41 AF; SSR+KD 81.10 ± 0.22 AA and 1.58 ± 0.18 AF. The same runs increased effective rank from 131.37 to 165.69 and reduced mean absolute off-diagonal prototype cosine from 0.335 to 0.064.

For capacity-style summaries, Cap@*θ* is the number of classes whose seen-class or task-incremental accuracy remains above threshold *θ*. Because threshold capacity depends on task order, the manuscript treats capacity as an operational proxy and emphasizes AA, AF, effective rank, and prototype overlap. Random-order controls are included to prevent overinterpreting TIL capacity.

CUB mask experiments used official CUB segmentation masks with frozen dense ResNet-18 features and a class-conditioned lightweight decoder. The strongest high-resolution setting used 192-pixel masks, seen-class SSR, *λ*_sp_ = 0.05, and *λ*_KD_ = 2.0, improving mIoU from 67.89 ± 0.34 to 71.58 ± 0.45 and Dice from 79.99 ± 0.27 to 82.73 ± 0.36.

### Transfer to camouflaged-object detection

The COD transfer experiment trained on COD10K-train and evaluated on CAMO, CHAMELEON, and COD10K-test (*54*). The decoder used channel-level SSR rather than class-prototype SSR because COD is a foreground-background segmentation problem with heavy texture ambiguity. Metrics included *S*_*α*_, weighted ***F***_*β*_, MAE, mIoU, and Dice. The compact paired summary is baseline 0.7440 and channel-level SSR 0.7572 for macro *S*_*α*_−MAE. This experiment is reported as supplementary transfer evidence; the main table reports the matched classification, editing, and adapter protocols.

### Transfer to sequential model-memory editing

KnowEdit-style edit streams tested whether SSR can regularize model-memory updates beyond image classifiers (*55, 56*). The lightweight editor is a fine-tuning-style update with SSR, spectral, and anchor terms applied to the edited weight subspace; true fine-tuning disables these regularizers. Evaluation followed the standard distinction between edit efficacy and locality. In ZsRE streams, tuned SSR-FT improved locality over true fine-tuning in 50-, 100-, 200-, 300-, and 400-edit settings; the 50-edit setting improved from 100.0/2.0 efficacy/locality for true fine-tuning to 100.0/13.0 for SSR-FT. WikiCounterFact and WikiRecent layer-17 matched controls likewise improved locality over true fine-tuning. These runs support a matched FT-style editing probe within the evaluated editor family. They are not intended to replace direct, same-budget comparisons with specialized editing systems such as ROME, MEMIT, MEND, WISE, or AlphaEdit (*57–62*).

### Additional LLM transfer

To test whether the GPT-2 XL locality trend extends beyond one checkpoint family, we evaluated additional Qwen and Llama models (*63–65*) on the same paired KnowEdit slices used for the main transfer analysis. The fixed grid kept the same two editors, the same three dataset families, and the same short edit budgets:

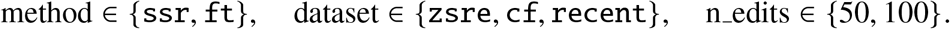

The additional checkpoint set was

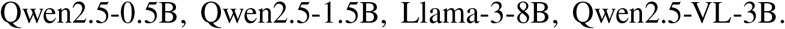

A comparison was included only when both efficacy and locality were available for FT and SSR-FT under the same model, dataset, and edit-budget setting. The additional-checkpoint editing claim in the manuscript rests on completed paired rows for Qwen2.5-VL-3B and Llama-3-8B. Across these completed pairs, SSR-FT retains ceiling efficacy or improves efficacy while increasing locality.

The GPT-2 XL control family establishes the main FT-style editing trend; Fig. S2 asks whether the same direction persists in paired additional-checkpoint slices.

**Table S3:**
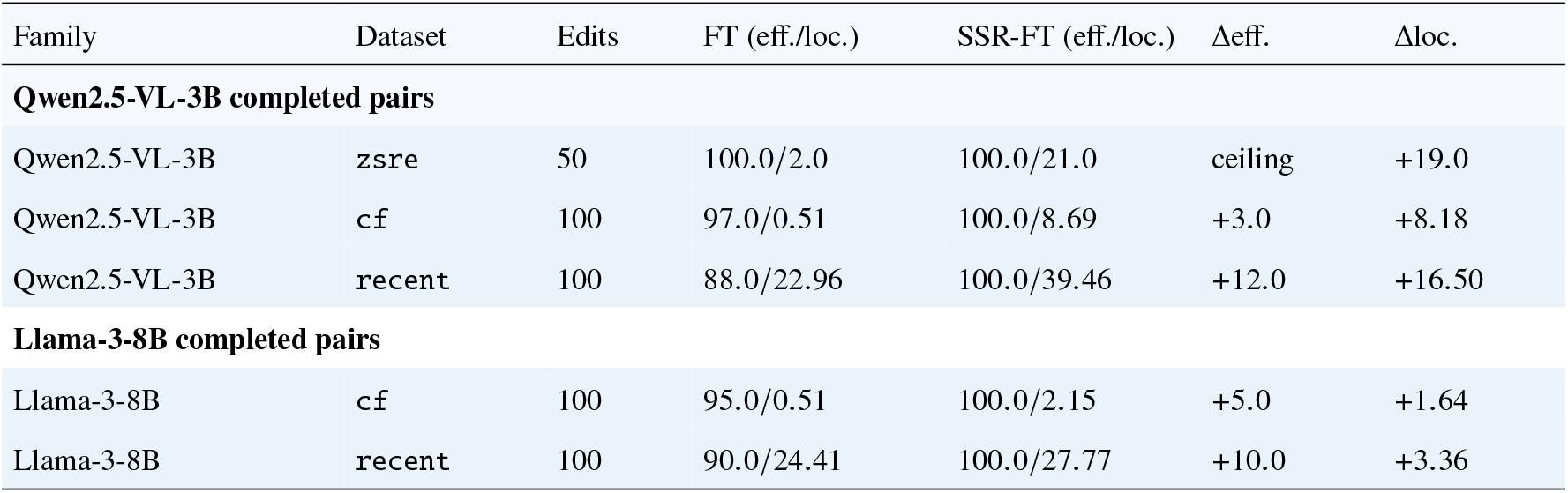
SSR-FT improves the locality–efficacy trade-off across completed additional-checkpoint KnowEdit pairs. Each row reports efficacy/locality for true fine-tuning and SSR-FT under the same model, dataset, and edit budget. In the completed Qwen2.5 and Llama pairs shown here, SSR-FT increases locality while retaining ceiling efficacy or improving efficacy.

#### Transfer to low-rank plastic subspaces

To test whether the same prior extends to low-rank plastic subspaces, we used the same CUB200 ResNet-18 feature representation but inserted a trainable rank-16 bottleneck adapter,

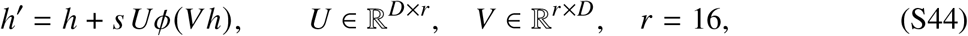

where *s* is a scale factor and *ϕ* is a nonlinear activation. SSR was applied to classifier prototypes and to the rows of ***U***, interpreted as output-basis directions of the adapter. The adapter-basis overlap reported in Table 2 is

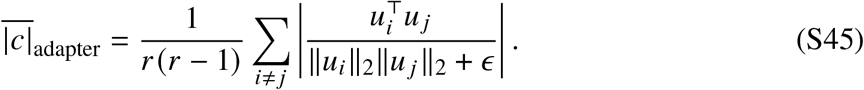

**Table 2:** SSR+KD improves the matched low-rank adapter accuracy–forgetting trade-off. Mean ± SEM over three seeds. Metric directions are shown in the headers. The blue row marks the adapter configuration used in the main text: SSR+KD improves the KD-only accuracy–forgetting trade-off and produces the strongest prototype geometry among the matched adapter variants.

| Method | AA ↑ | AF ↓ | Eff. rank ↑ | Proto. \|cos\| ↓ |
| --- | --- | --- | --- | --- |
| Baseline | 76.69±0.03 | 4.14±0.06 | 154.87±0.29 | 0.101±0.002 |
| KD | 79.26±0.16 | 3.01±0.19 | 160.15±0.26 | 0.101±0.001 |
| SSR | 76.98±0.86 | 4.16±0.86 | 161.58±0.06 | 0.084±0.001 |
| SSR+KD | 79.78±0.20 | 2.58±0.22 | 168.08±0.21 | 0.075±0.000 |

The experiment used three seeds and 20 epochs per task. Within this matched frozen-feature adapter protocol, SSR+KD gave the strongest accuracy–forgetting trade-off among the tested adapter variants; standard full LoRA systems are treated as a separate benchmark family rather than a matched comparator (*36*).

**Figure S2:**
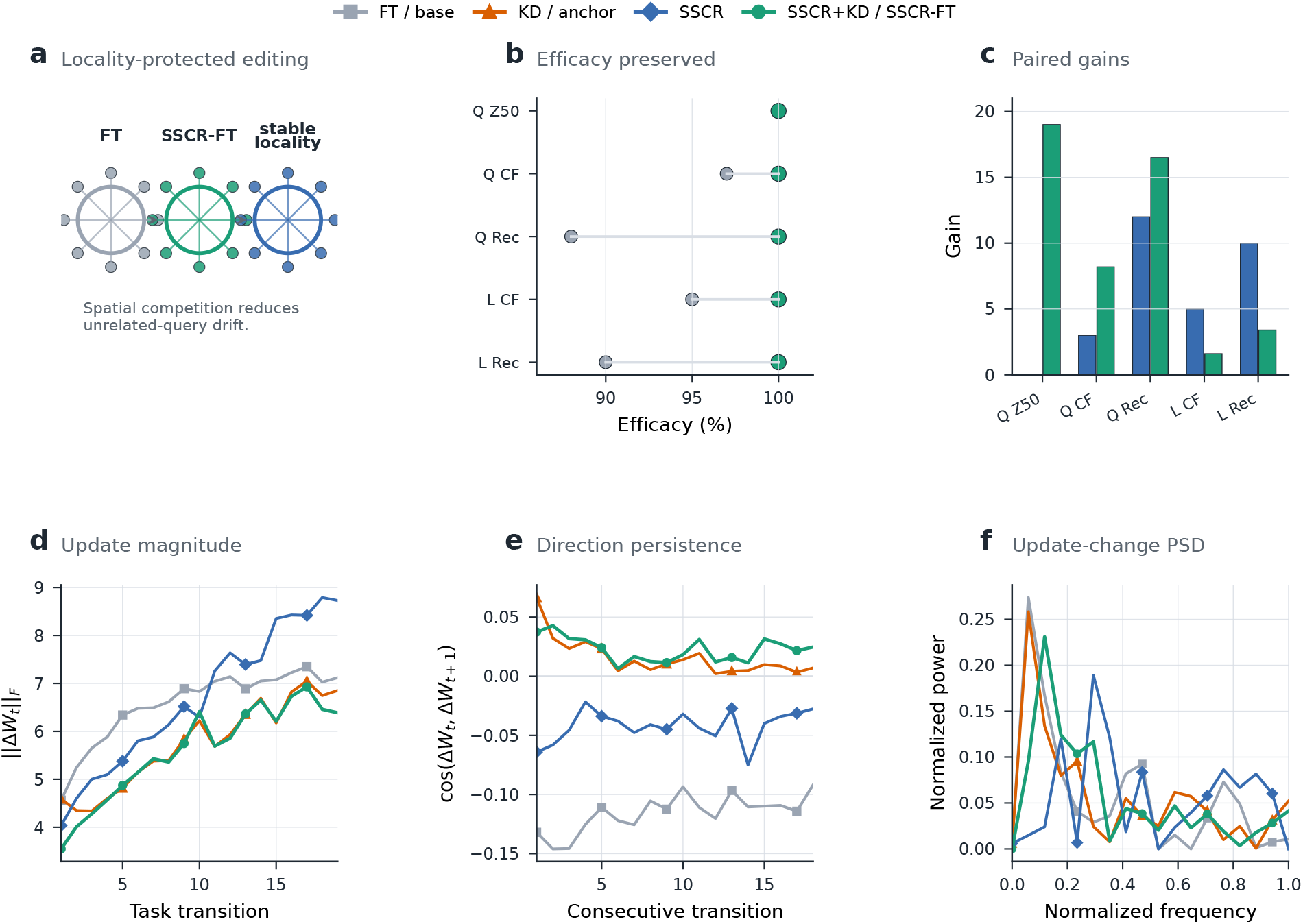
Additional-checkpoint editing support and task-resolution dynamics. **a**, Locality-protected sequential-editing schematic. **b**, Paired efficacy readouts for Qwen2.5-VL-3B and Llama-3-8B slices. **c**, Paired Δefficacy and Δlocality summary bars for the same slices. **d**, Task-resolution update magnitude is plotted from 19 CUB200 classifier-weight transitions saved at task boundaries. **e**, Consecutive update-direction cosine is plotted for each adjacent task transition. **f**, Update-direction-change PSD is computed from the same transition series and normalized to total spectral power.

### Task-resolution dynamics

To test whether SSR changes learning dynamics over task time rather than only the internal structure of one update matrix, we analyzed CUB200 classifier checkpoints 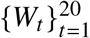 saved after each task and formed task-transition updates Δ*W*_*t*_ = *W*_*t*_ − *W*_*t*−1_. From these we constructed scalar time series including update magnitude

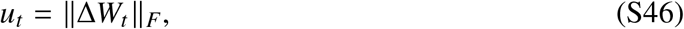

and update-direction continuity

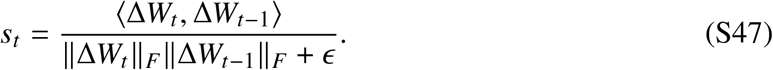

For scalar series *x*_*t*_, the normalized autocorrelation was

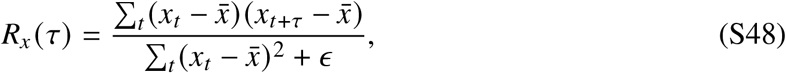

and the corresponding one-sided power spectrum was obtained from the Fourier transform of the mirrored non-negative-lag autocorrelation, following the Wiener–Khinchin relation. We also directly computed lagged cosine autocorrelation between update vectors,

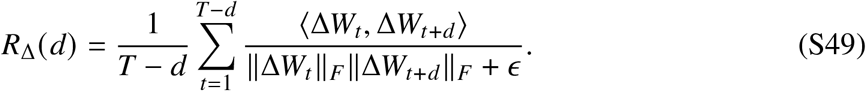

result should be interpreted as a task-resolution temporal analysis rather than an epoch- or step-level dynamics analysis. In this setting, KD-only already reduced mean update norm, increased lag-1 update cosine, and lowered the high-frequency fraction of the update-autocorrelation spectrum relative to the baseline. SSR+KD gave a further small but consistent improvement on these three temporal-dynamics metrics.

The task-resolution dynamics in Fig. S2 make the support explicit: KD-only already smooths the update stream, whereas SSR+KD gives the strongest combined temporal profile in this task-transition analysis.

### Axis-aware Fourier scope analysis

This analysis answers a different question from the temporal one above. For task transition *t*, the classifier update is Δ*W*_*t*_ = *W*_*t*_ −*W*_*t*−1_. The flattened FFT statistic computes

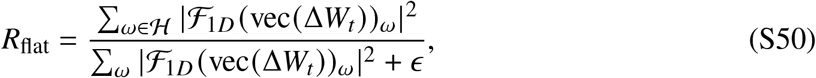

where *ℋ* is the high-frequency band. Because vectorization imposes an arbitrary order, we also used class-axis, feature-axis, and two-dimensional radial spectra,

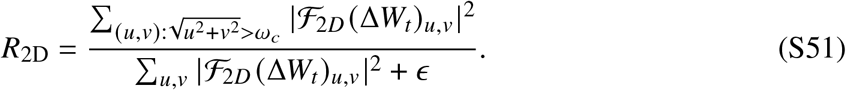

Both ***U***_flat_ and ***U***_2D_ are relative quantities: the numerator is the high-frequency band power and the denominator is the total spectral power under the same transform. We also recorded the corresponding absolute high-frequency band power,

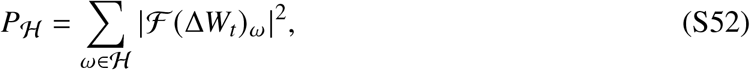

using the same band definition as the ratio. Permutation-null controls shuffled entries of Δ*W*_*t*_ before the flattened FFT. SSR+KD reduced update norm in CUB task checkpoints and reorganized axis-aware normalized high-frequency ratios; the same-convention absolute high-frequency band powers were also recorded to avoid interpreting ratio changes alone. Therefore, the Fourier panels define the scope of the mechanism: the primary support is high-rank, low-overlap geometry, localized CUB attention, and task-level temporal persistence.

The Fourier views in Fig. S3 summarize how SSR changes the matched CUB runs through representational allocation, localized evidence, and persistent task-level updates under axis-aware and permutation-null controls.

### Supplementary figure support

Figure S3 summarizes supplementary analyses with exact values provided in Tables S4–S7. The Split-CIFAR-100 trajectory and endpoint summaries show that the matched trade-off shift in Fig. 4 is sustained across the full task stream rather than arising from a favorable endpoint average alone. The CUB order-robustness, rank-retention, and Fourier panels separate three mechanistic claims: the geometry shift is robust to task ordering, KD converts that geometry into retention, and the same spatial prior reorganizes axis-aware spectra while preserving the high-rank, low-overlap mechanism.

### Computational reproducibility

Training configurations, seed lists, run summaries, and figure-generation records were retained for the artificial-network experiments. Dataset inputs were obtained from the public sources cited above. The reported numerical summaries were generated from saved per-seed results or task-boundary checkpoints using the same metric definitions described in this supplement.

**Figure S3:**
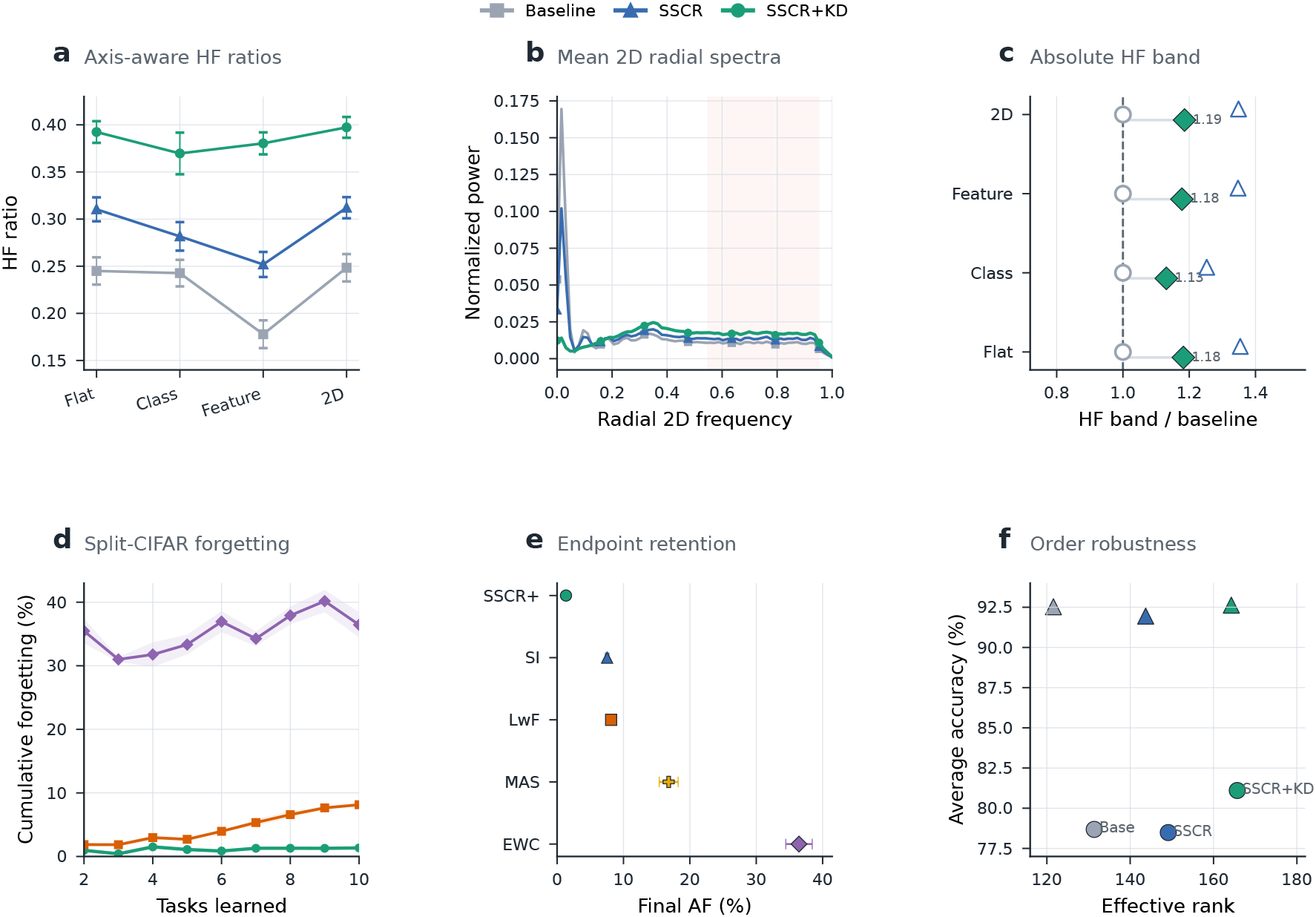
Combined supplementary atlas for Fourier scope analysis, retention progression, and geometry transfer. **a**, Normalized high-frequency ratios are computed from the CUB classifier-update Fourier analysis. **b**, Mean radial two-dimensional spectra are plotted from the same task-transition spectra, with HF-band shading. **c**, Absolute high-frequency band powers are computed from per-view high-ratio and total-power summaries. **d**, Split-CIFAR-100 forgetting trajectories are computed from seed-level accuracy matrices. **e**, Final Split-CIFAR-100 forgetting endpoints are computed from the same seed-level matrices. **f**, CUB200 semantic- and random-order rank– accuracy scatter summarizes the matched classification runs. Tables S4–S7 provide exact values.

**Table S4:** SSR+KD couples higher-rank geometry to the strongest CUB200 retention trade-off. SSR+KD combines the largest effective-rank expansion, the smallest mean prototype overlap, and the strongest average-accuracy / forgetting trade-off.

| Method | AA ↑ | AF ↓ | Effective rank ↑ | Mean $\vert \cos\vert \downarrow$ |
| --- | --- | --- | --- | --- |
| Baseline | 78.68±0.39 | 3.49±0.41 | 131.37 | 0.335 |
| SSR | 78.49±0.19 | 3.54±0.32 | 149.14 | 0.095 |
| SSR+KD | 81.10±0.22 | 1.58±0.18 | 165.69 | 0.064 |

**Table S5:** SSR+KD combines the lowest update magnitude with the strongest temporal-persistence profile. The columns are task-resolution metrics aligned with the main mechanism claim: lower mean update norm, more positive adjacent-update cosine, and lower high-frequency autocorrelation power. KD-only already smooths the update stream relative to the baseline, whereas SSR+KD gives the strongest profile on all three metrics.

| Method | mean $\ \Delta W\ _F \downarrow$ | lag-1 cosine ↑ | PSD high ↓ |
| --- | --- | --- | --- |
| Baseline | 6.575 | -0.116 | 0.602 |
| KD | 5.736 | 0.016 | 0.526 |
| SSR | 6.697 | -0.042 | 0.561 |
| SSR+KD | **5.613** | **0.023** | **0.520** |

**Table S6:** SSR+KD lowers update norm and redistributes axis-aware high-frequency ratios. HF values are normalized high-frequency ratios, i.e. high-frequency band power divided by total spectral power. This analysis scopes the mechanism around representational allocation, localized evidence, and task-level temporal persistence.

| Method | $\ \Delta W\ _2 \downarrow$ | Flat HF | Class HF | Feature HF | 2D HF |
| --- | --- | --- | --- | --- | --- |
| Baseline | 6.575±0.726 | 0.245±0.014 | 0.243±0.014 | 0.178±0.015 | 0.248±0.015 |
| SSR | 6.697±1.453 | 0.310±0.013 | 0.282±0.015 | 0.252±0.013 | 0.312±0.011 |
| SSR+KD | 5.613±0.962 | 0.392±0.011 | 0.370±0.022 | 0.380±0.012 | 0.397±0.011 |

**Table S7:** Absolute high-frequency band powers under the same Fourier controls. Values are means over task transitions, computed as normalized HF ratio times total spectral power for each transform view. These same-convention absolute band powers are reported alongside the ratios so the Fourier scope analysis is not interpreted from relative normalization alone.

| Method | Flat $P_H$ | Class $P_H$ | Feature $P_H$ | 2D $P_H$ |
| --- | --- | --- | --- | --- |
| Baseline | 5.52$\times 10^5$ | 2.10 | 13.93 | 1.11$\times 10^6$ |
| SSR | 7.47$\times 10^5$ | 2.63 | 18.76 | 1.50$\times 10^6$ |
| SSR+KD | 6.52$\times 10^5$ | 2.37 | 16.41 | 1.32$\times 10^6$ |

## Notes

https://github.com/ydchen0806/ssr-ai-reproducibility

### Competing Interest Statement

The authors have declared no competing interest.

## References and Notes

1. D. O. Hebb, The Organization of Behavior: A Neuropsychological Theory (Wiley, New York) (1949).

2. T. V. P. Bliss, T. Lomo, Long-lasting potentiation of synaptic transmission in the dentate area of the anaesthetized rabbit following stimulation of the perforant path. The Journal of Physiology 232 (2), 331–356 (1973), doi:10.1113/jphysiol.1973.sp010273.

3. H. Markram, J. Lubke, M. Frotscher, B. Sakmann, Regulation of synaptic efficacy by coincidence of postsynaptic APs and EPSPs. Science 275 (5297), 213–215 (1997), doi: 10.1126/science.275.5297.213.

4. G.-q. Bi, M.-m. Poo, Synaptic modifications in cultured hippocampal neurons: dependence on spike timing, synaptic strength, and postsynaptic cell type. The Journal of Neuroscience 18 (24), 10464–10472 (1998), doi:10.1523/JNEUROSCI.18-24-10464.1998.

5. E. Oja, Simplified neuron model as a principal component analyzer. Journal of Mathematical Biology 15 (3), 267–273 (1982), doi:10.1007/BF00275687.

6. E. L. Bienenstock, L. N. Cooper, P. W. Munro, Theory for the development of neuron selectivity: orientation specificity and binocular interaction in visual cortex. The Journal of Neuroscience 2 (1), 32–48 (1982), doi:10.1523/JNEUROSCI.02-01-00032.1982.

7. G. G. Turrigiano, S. B. Nelson, Homeostatic plasticity in the developing nervous system. Nature Reviews Neuroscience 5 (2), 97–107 (2004), doi:10.1038/nrn1327.

8. Clopath, L. Busing, E. Vasilaki, W. Gerstner, Connectivity reflects coding: a model of voltage-based STDP with homeostasis. Nature Neuroscience 13 (3), 344–352 (2010), doi: 10.1038/nn.2479.

9. L. F. Abbott, S. B. Nelson, Synaptic plasticity: taming the beast. Nature Neuroscience 3 (11s), 1178–1183 (2000), doi:10.1038/81453.

10. J. C. Magee, Dendritic integration of excitatory synaptic input. Nature Reviews Neuroscience 1 (3), 181–190 (2000), doi:10.1038/35044552.

11. L. Sabatini, T. G. Oertner, K. Svoboda, The life cycle of Ca2+ ions in dendritic spines. Neuron 33 (3), 439–452 (2002), doi:10.1016/S0896-6273(02)00573-1.

12. M. Matsuzaki, N. Honkura, G. C. R. Ellis-Davies, H. Kasai, Structural basis of long-term potentiation in single dendritic spines. Nature 429 (6993), 761–766 (2004), doi:10.1038/nature02617.

13. D. Harvey, K. Svoboda, Locally dynamic synaptic learning rules in pyramidal neuron dendrites. Nature 450 (7173), 1195–1200 (2007), doi:10.1038/nature06416.

14. Holtmaat, K. Svoboda, Experience-dependent structural synaptic plasticity in the mammalian brain. Nature Reviews Neuroscience 10 (9), 647–658 (2009), doi:10.1038/nrn2699.

15. T. Branco, B. A. Clark, M. Hausser, Dendritic discrimination of temporal input sequences in cortical neurons. Science 329 (5999), 1671–1675 (2010), doi:10.1126/science.1189664.

16. R. Yuste, Dendritic spines and distributed circuits. Neuron 71 (5), 772–781 (2011), doi:10.1016/j.neuron.2011.07.024.

17. W. Denk, H. Horstmann, Serial block-face scanning electron microscopy to reconstruct three-dimensional tissue nanostructure. PLoS Biology 2 (11), e329 (2004), doi:10.1371/journal.pbio.0020329.

18. Motta, et al., Dense connectomic reconstruction in layer 4 of the somatosensory cortex. Science 366 (6469), eaay3134 (2019), doi:10.1126/science.aay3134.

19. MICrONS Consortium, et al., Functional connectomics spanning multiple areas of mouse visual cortex. bioRxiv (2021), doi:10.1101/2021.07.28.454025.

20. N. L. Turner, et al., Reconstruction of neocortex: Organelles, compartments, cells, circuits, and activity. Cell 185 (6), 1082–1100 (2022), doi:10.1016/j.cell.2022.01.023.

21. Shapson-Coe, et al., A petavoxel fragment of human cerebral cortex reconstructed at nanoscale resolution. Science 384 (6696), eadk4858 (2024), doi:10.1126/science.adk4858.

22. J. I. Arellano, R. Benavides-Piccione, J. DeFelipe, R. Yuste, Ultrastructure of dendritic spines: correlation between synaptic and spine morphologies. Frontiers in Neuroscience 1, 131–143 (2007), doi:10.3389/neuro.01.1.1.010.2007.

23. T. M. Bartol, et al., Nanoconnectomic upper bound on the variability of synaptic plasticity. eLife 4, e10778 (2015), doi:10.7554/eLife.10778.

24. S. Dorkenwald, et al., Binary and analog variation of synapses between cortical pyramidal neurons. eLife 11, e76120 (2022), doi:10.7554/eLife.76120.

25. S. Fusi, P. J. Drew, L. F. Abbott, Cascade models of synaptically stored memories. Neuron 45 (4), 599–611 (2005), doi:10.1016/j.neuron.2005.02.001.

26. L. Bloodgood, B. L. Sabatini, Neuronal activity regulates diffusion across the neck of dendritic spines. Science 310 (5749), 866–869 (2005), doi:10.1126/science.1114816.

27. T. Kleindienst, J. Winnubst, C. Roth-Alpermann, T. Bonhoeffer, M. E. Larkum, Activity-dependent clustering of functional synaptic inputs on developing hippocampal dendrites. Neuron 72 (6), 1012–1024 (2011), doi:10.1016/j.neuron.2011.10.015.

28. Olshausen, D. J. Field, Emergence of simple cell receptive fields by learning a sparse code for natural images. Nature 381 (6583), 607–609 (1996), doi:10.1038/381607a0.

29. N. C. Rust, J. J. DiCarlo, Selectivity and tolerance (“invariance”) both increase as visual information propagates from cortical area V4 to IT. Journal of Neuroscience 30 (39), 12978–12995 (2010), doi:10.1523/JNEUROSCI.0179-10.2010.

30. T. K. Hensch, Critical period plasticity in local cortical circuits. Nature Reviews Neuroscience 6 (11), 877–888 (2005), doi:10.1038/nrn1787.

31. N. Takahashi, et al., Locally synchronized synaptic inputs. Science 335 (6066), 353–356 (2012), doi:10.1126/science.1210362.

32. Wilson, D. E. Whitney, B. Scholl, D. Fitzpatrick, Orientation selectivity and the functional clustering of synaptic inputs in primary visual cortex. Nature Neuroscience 19 (8), 1003–1009 (2016), doi:10.1038/nn.4323.

33. M. F. Iacaruso, I. T. Gasler, S. B. Hofer, Synaptic organization of visual space in primary visual cortex. Nature 547 (7664), 449–452 (2017), doi:10.1038/nature23019.

34. K. He, X. Zhang, S. Ren, J. Sun, Deep residual learning for image recognition, in Proceedings of the IEEE Conference on Computer Vision and Pattern Recognition (2016), pp. 770–778.

35. Wah, S. Branson, P. Welinder, P. Perona, S. Belongie, The Caltech-UCSD Birds-200-2011 Dataset, Tech. Rep. CNS-TR-2011-001, California Institute of Technology (2011).

36. J. Hu, et al., LoRA: Low-rank adaptation of large language models, in International Conference on Learning Representations (2022).

37. Govindarajan, I. Israely, S.-Y. Huang, S. Tonegawa, The dendritic branch is the preferred integrative unit for protein synthesis-dependent LTP. Neuron 69 (1), 132–146 (2011), doi: 10.1016/j.neuron.2010.12.008.

38. Hinton, O. Vinyals, J. Dean, Distilling the knowledge in a neural network, in NIPS Deep Learning and Representation Learning Workshop (2015).

39. Z. Li, D. Hoiem, Learning without forgetting. IEEE Transactions on Pattern Analysis and Machine Intelligence 40 (12), 2935–2947 (2018), doi:10.1109/TPAMI.2017.2773081.

40. R. Hadsell, D. Rao, A. A. Rusu, R. Pascanu, Embracing change: continual learning in deep neural networks. Trends in Cognitive Sciences 24 (12), 1028–1040 (2020), doi:10.1016/j.tics.2020.09.004.

41. M. van de Ven, H. T. Siegelmann, A. S. Tolias, Brain-inspired replay for continual learning with artificial neural networks. Nature Communications 11 (1), 4069 (2020), doi:10.1038/s41467-020-17866-2.

42. M. De Lange, et al., A continual learning survey: Defying forgetting in classification tasks. IEEE Transactions on Pattern Analysis and Machine Intelligence 44 (7), 3366–3385 (2021), doi:10.1109/TPAMI.2021.3057446.

43. L. Wang, X. Zhang, H. Su, J. Zhu, A comprehensive survey of continual learning: Theory, method and application. IEEE Transactions on Pattern Analysis and Machine Intelligence 46 (8), 5362–5383 (2024), doi:10.1109/TPAMI.2024.3367329.

44. Chaudhry, P. K. Dokania, T. Ajanthan, P. H. S. Torr, Riemannian walk for incremental learning: Understanding forgetting and intransigence, in European Conference on Computer Vision (2018), pp. 532–547.

45. N. Diaz-Rodriguez, V. Lomonaco, D. Filliat, D. Maltoni, Don’t forget, there is more than forgetting: new metrics for continual learning, in NeurIPS Workshop on Continual Learning (2018).

46. Krizhevsky, G. Hinton, Learning multiple layers of features from tiny images, Tech. rep., University of Toronto (2009).

47. Y. Le, X. Yang, Tiny ImageNet Visual Recognition Challenge. CS 231N Course Report, Stanford University (2015).

48. J. Kirkpatrick, et al., Overcoming catastrophic forgetting in neural networks. Proceedings of the National Academy of Sciences 114 (13), 3521–3526 (2017), doi:10.1073/pnas.1611835114.

49. Zenke, B. Poole, S. Ganguli, Continual learning through synaptic intelligence, in Proceedings of the 34th International Conference on Machine Learning (2017), pp. 3987–3995.

50. R. Aljundi, F. Babiloni, M. Elhoseiny, M. Rohrbach, T. Tuytelaars, Memory aware synapses: Learning what (not) to forget, in Proceedings of the European Conference on Computer Vision (2018), pp. 139–154.

51. S.-A. Rebuffi, A. Kolesnikov, G. Sperl, C. H. Lampert, iCaRL: Incremental classifier and representation learning, in Proceedings of the IEEE Conference on Computer Vision and Pattern Recognition (2017), pp. 2001–2010.

52. Lopez-Paz, M. Ranzato, Gradient episodic memory for continual learning, in Advances in Neural Information Processing Systems, vol. 30 (2017).

53. P. Buzzega, M. Boschini, A. Porrello, S. Calderara, Dark experience for general continual learning: a strong, simple baseline, in Advances in Neural Information Processing Systems, vol. 33 (2020), pp. 15920–15930.

54. D.-P. Fan, et al., Camouflaged object detection. Proceedings of the IEEE/CVF Conference on Computer Vision and Pattern Recognition pp. 2777–2787 (2020).

55. N. Zhang, et al., A comprehensive study of knowledge editing for large language models. arXiv preprint arXiv:2401.01286 (2024).

56. P. Wang, et al., EasyEdit: An easy-to-use knowledge editing framework for large language models. arXiv preprint arXiv:2308.07269 (2024).

57. K. Meng, D. Bau, A. Andonian, Y. Belinkov, Locating and editing factual associations in GPT, in Advances in Neural Information Processing Systems, vol. 35 (2022), pp. 17359–17372.

58. K. Meng, A. S. Sharma, A. Andonian, Y. Belinkov, D. Bau, Mass-editing memory in a transformer, in International Conference on Learning Representations (2023).

59. Mitchell, C. Lin, A. Bosselut, C. Finn, C. D. Manning, Fast model editing at scale, in International Conference on Learning Representations (2022).

60. Mitchell, C. Lin, A. Bosselut, C. D. Manning, C. Finn, Memory-based model editing at scale, in International Conference on Machine Learning (2022).

61. J. Fang, et al., AlphaEdit: Null-space constrained knowledge editing for language models, in International Conference on Learning Representations (2025).

62. P. Wang, et al., WISE: Rethinking the knowledge memory for lifelong model editing of large language models, in Advances in Neural Information Processing Systems (2024).

63. Qwen Team, Qwen2.5 Technical Report. arXiv preprint arXiv:2412.15115 (2024), https://arxiv.org/abs/2412.15115.

64. Qwen Team, Qwen2.5-VL Technical Report. arXiv preprint arXiv:2502.13923 (2025), https://arxiv.org/abs/2502.13923.

65. Dubey, et al., The Llama 3 Herd of Models. arXiv preprint arXiv:2407.21783 (2024), https://arxiv.org/abs/2407.21783.

